# Adolescent alcohol exposure alters age-related progression of behavioral and neurotrophic dysfunction in the TgF344-AD model in a sex-specific manner

**DOI:** 10.1101/2024.07.17.603911

**Authors:** Nicole L. Reitz, Polliana T. Nunes, Lisa M. Savage

**Affiliations:** Department of Psychology, Binghamton University, State University of New York.

**Keywords:** Alzheimer’s disease, rat model, adolescent alcohol, hippocampus, neurotrophins

## Abstract

Alzheimer’s Disease (AD) and heavy alcohol use are widely prevalent and lead to brain pathology. Both alcohol-related brain damage (ABRD) and AD result in cholinergic dysfunction, reductions in hippocampal neurogenesis, and the emergence of hippocampal-dependent cognitive impairments. It is still unknown how ARBD caused during a critical developmental timepoint, such as adolescence, interacts with AD-related pathologies to accelerate disease progression later in life. The current study utilized a longitudinal design to characterize behavioral and pathological changes in a transgenic rat model of AD (TgF344-AD) following adolescent intermittent ethanol (AIE) exposure. We found that AIE accelerates cognitive decline associated with AD transgenes in female rats at 6 months of age, and male AD-rats are impaired on spatial navigation by 3-months with no additional deficits due to AIE exposure. Protein levels of various AD-pathological markers were analyzed in the dorsal and ventral hippocampus of male and female rats. The data suggests that AIE-induced alterations of the tropomyosin-related kinase A receptor (TrkA) / p75 neurotrophin receptor (p75NTR) ratio creates a brain that is vulnerable to age- and AD-related pathologies, which leads to an acceleration of cognitive decline, particularly in female rats.

## 1. Introduction

Chronic and heavy alcohol consumption is a risk factor for the age-related development of dementia, including Alzheimer’s disease (AD; León et al., 2021). Despite a growing literature on alcohol-AD interactions, sex differences have been insufficiently explored. Many studies have not determined whether sex differences exist in alcohol-AD interactions, despite the fact that women are more likely to develop AD and experience a more rapid cognitive decline and brain atrophy progression upon AD diagnosis (Holland et al., 2013; Lin et al., 2015; Mielke, 2018). In addition, there is evidence that female rodents are more sensitive to alcohol-related brain damage (Kipp et al., 2021; Maynard et al., 2018). Emerging evidence suggests that female rodents may be more vulnerable to alcohol-induced acceleration of AD-like pathology (Barnett et al., 2022; Tucker et al., 2022).

Furthermore, many alcohol and AD interaction studies focus solely on alcohol use disorder (AUD) in adulthood (Sanna et al., 2023), leaving a knowledge gap regarding the impact of developmental alcohol exposure on AD symptom and pathology progression. Thus, the effects of aging following developmental alcohol exposure on AD-related behavioral dysfunction and pathology remains underexplored. This is somewhat surprising given that rodent models have revealed that adolescent ethanol exposure presents unique pathologies, some similar to pathologies seen in AD and advanced aging, such as altered neuroimmune signaling, cholinergic dysfunction, and reduced hippocampal neurogenesis (Coleman et al., 2011, 2021; Vetreno et al., 2014). Thus, a history of heavy alcohol use during adolescence may prime the brain for AD-related pathologies during the aging process.

Rodent models that display age-related progression of AD-related pathologies (amyloid plaque and tau expression) and behavioral dysfunction are key to understanding AD pathogenesis, risk factors, and novel therapeutic approaches (Drummond & Wisniewski, 2017). One model that has well characterized AD-neuropathologies that align with progressive cognitive decline across the lifespan is the TgF344-AD rat model (Cohen et al., 2013). At 3 months of age, TgF344-AD rats (AD-rats) perform significantly worse than wildtype (WT) controls on a ventral hippocampus-dependent social interaction task (Chaney et al., 2021). By 5 months of age, before Aβ plaques form, AD male rats take longer to meet criteria on a delayed-non-match-to-sample task, although percentage correct remains consistent at each delay (Muñoz-Moreno et al., 2018). The Morris Water Maze (MWM), a standard test for hippocampus-dependent spatial memory, reveals deficits in reversal learning as early as 6 months old, persisting until 16 months (Berkowitz et al., 2018; Rorabaugh et al., 2017; Voorhees et al., 2019). Interestingly, the lack of initial learning deficits on the MWM suggests their ability to perform tasks reliant solely on the hippocampus (initial spatial learning) remain intact in adulthood. However, early deficits emerge in cognitive flexibility, such as reversal learning, that requires communication between the hippocampus, basal forebrain, and cortical regions. Novel object recognition (NOR), dependent on the interactions between the hippocampus and entorhinal cortex, reveals intact memory at 3 and 6 months in both AD male and female rats, but an inability to discriminate between objects from 9 to 12 months (Chaney et al., 2021; Galloway et al., 2018). In female rats, novel object location/place (NOP), a neurogenesis-dependent task, displays impairment at 9-12 months, but not at 5-8 months (Galloway et al., 2018). However, these studies did not use sex as a factor in the statistical analyses to determine unique sex-specific differences.

Exploring sex differences in cognitive performance in this model has been limited. The delayed rewarded T-maze alternation shows declining performance at 12 and 18 months in male AD rats, but only at 18 months in AD female rats (Saré et al., 2020). Similarly, in the MWM, deficits emerge at 9 and 12 months in AD male rats, while AD females exhibit impairments only at 12 months of age (Srivastava et al., 2023). Alternatively, at 9 months, female AD rats display increased anxiety, not observed in males (Srivastava et al., 2023). Sex differences in other cognitive tasks in this AD model have not been fully characterized. Thus, AD male rats appear to display spatial memory impairment earlier than their female counterparts, but female AD rats selectively display increased anxiety with advanced age.

However, the impact of developmental ethanol exposure on cognitive decline trajectory in the AD model, and many other models, remains unexplored. Adolescent intermittent ethanol exposure (AIE) in wildtype (WT) rodents reduces cholinergic markers in the medial septum/diagonal band and neurogenesis in the hippocampus, increasing susceptible to early age-related cognitive decline (Fernandez & Savage, 2017; Reitz et al., 2021). The loss of cholinergic expression in neurons within the basal forebrain results from AIE-induced epigenetic suppression of genes driving this phenotype (Vetreno et al., 2020). Cholinergic neurons in the basal forebrain rely on retrograde transport of nerve growth factor (NGF) for maintenance (Cuello et al., 2007; Hagg et al., 1988, 1989), and NGF levels are affected by AIE exposure (Vetreno et al., 2018) and in AD (Cuello et al., 2019; Scott & Crutcher, 1994). AIE exposure alone suppresses gene markers for TrkA (high-affinity NGF receptor) and p75 (low-affinity NGF receptor) in the basal forebrain (Vetreno et al., 2020), and reduces choline acetyltransferase (ChAT) protein levels in the basal forebrain and vesicular acetylcholine transporter (VAChT) protein levels in the orbital frontal cortex (Kipp et al., 2021). Similar patterns are observed in human AD, such as decreased VAChT in CA1, CA3, and prefrontal cortex (basal forebrain projection sites) and decreased TrkA in the basal forebrain (Gilmor et al., 1999; Ginsberg et al., 2019; Parent et al., 2013). Furthermore, ethanol exposure during adolescences does accelerate adult brain Aβ_42_protein levels in female 3xTg-AD mouse model (Barnett et al., 2022) and the APP (amyloid precursor protein)/PSEN (presenilin) mouse model (Ledesma et al., 2021).

Therefore, studies are crucial to determining whether/when sex differences emerge during aging following adolescent ethanol exposure in rodent models, particularly in AD models. AIE-induced changes across the dorsal and ventral hippocampus may render rodents more vulnerable to aging-related cognitive decline, potentially accelerating cognitive decline in the presence of genetic mutations like APP and presenilin 1 (PS1). The dorsal and ventral sectors of the hippocampus were examined as rodents with AD transgenes, the plaque load in the ventral hippocampus exceeds that observed in the dorsal hippocampus (Russo et al., 2021; Tsui et al., 2022).

Given prior findings of heightened sensitivity to AIE-induced effects in 3x-Tg AD female mice (Barnett et al, 2022; Ledesma et al, 2021), and the growing human literature on gender differences in AD development and progression, female AD rodents may exhibit increased vulnerability to AD-like pathology and accelerated cognitive decline following adolescent ethanol exposure compared to male AD-rats. These sex-specific hypotheses were tested in a unique longitudinal design using standard behavioral protocols for spatial memory (plus maze spontaneous alternation and operant delayed nonmatching to position [DNMTP]) and social recognition memory at three age ranges (3, 6, and 9 months) with brain samples collected at 11 months for analysis of alterations of neurotrophin-related and neuropathological protein levels.

## 2. Materials and Methods

### 2.1 Subjects

Male (*N* = 48) and female (*N* = 48) rats were bred at Binghamton University using female wild-type (WT) F344 rats (obtained from Charles River, Stoneridge, NY) and male TgF344-AD rats (obtained from RRRC, Columbia, MO). Of the 96 total rats, 48 had AD transgenes (Cohen et al., 2013) and 48 were their WT littermates. On postnatal day (P) 21, all rats were weaned from the dam and genotyped using ear-punched tissue. Tissue was sent to Transnetyx (Cordova, TN) for automated genotyping. All rats ranged from P27-29 at the start of ethanol exposure (see below). Only one rat per sex per litter was randomly assigned to a given exposure group. Rats were pair-housed post-weaning and given free access to food and water. All rats were housed in a temperature-controlled colony room that maintains a 12-hour light/dark cycle from 7:00AM to 7:00PM. Rats were given wood chew blocks and crinkle paper as enrichment and were housed in standard bedding. All procedures and housing are in accordance with the Institutional Animal Care and Use Committee at Binghamton University.

### 2.2 Adolescent Intermittent Ethanol Exposure

Rats of both sexes and genotypes were exposed to either a binge-type forced gavage ethanol exposure (AIE: *N* = 48) or tap water control gavage exposure (CON: *N* = 48). Rats were exposed to 16 intragastric gavages of either 20% ethanol (*v/v*) or tap water, both of which were administered at a dose of 5g/kg (see *Figure 1*). The exposure cycle followed a two-day on/off cycle from P28-58. Tail blood samples were collected one hour after the eighth exposure (P41) to ensure binge levels of ethanol consumption (BEC > 0.08) were met. All blood samples were processed using an AM1 Alcohol Analyzer (ANALOX Instruments, England) and the blood ethanol content (BEC) of each animal was recorded. Following the final ethanol or water exposure, all rats remained in an abstinence (no ethanol) phase for the remaining duration of the study.

**Figure 1.**
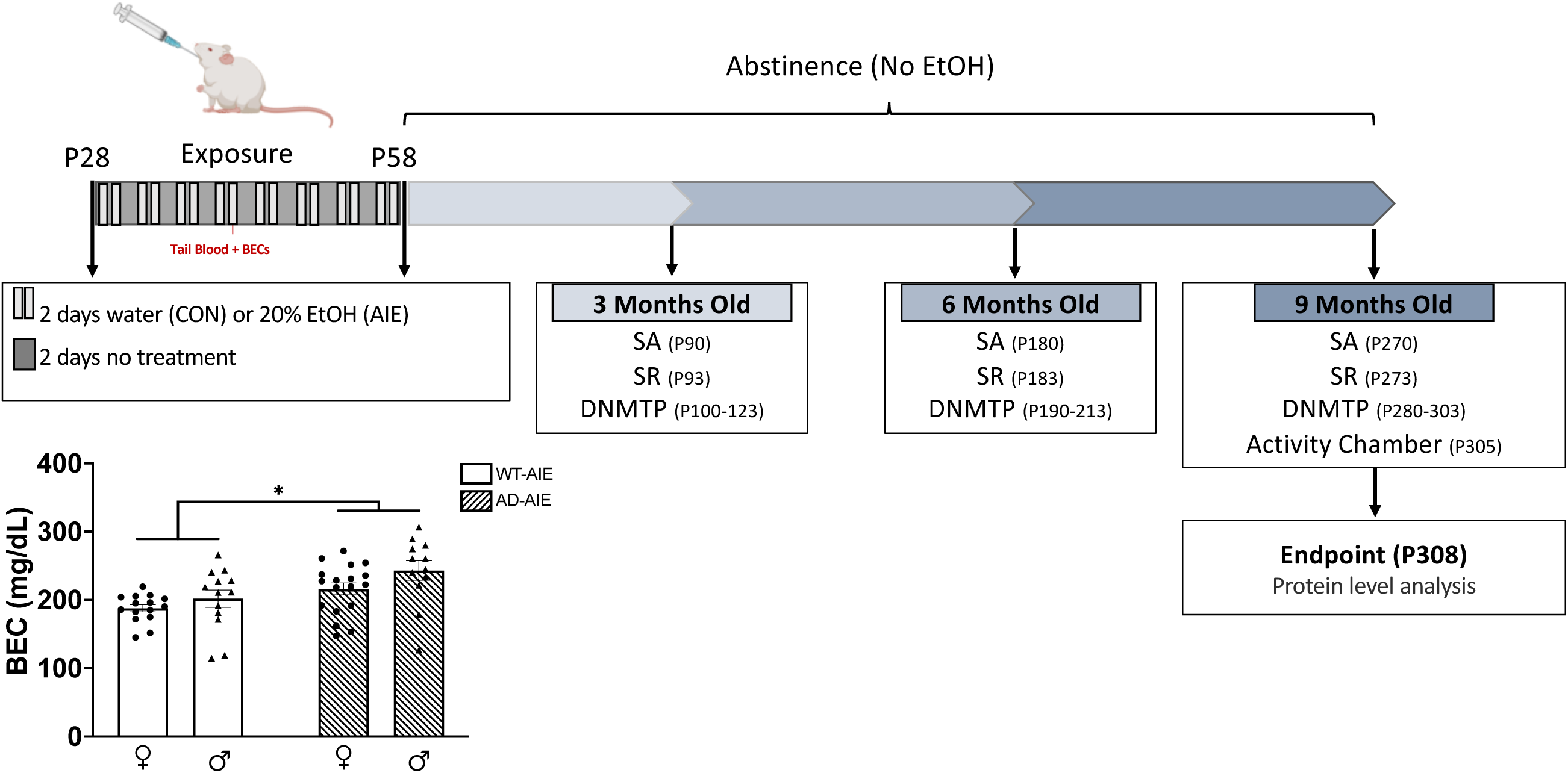
Schematic of experimental design. All rats underwent a gavage procedure of 20% ethanol (AIE) or tap water (CON) for two days on/two days off from postnatal day (P) 28-58. Blood ethanol concentrations (BECs) exceeded binge levels (>80mg/dL). BECs were greater for TgF344-AD rats (dashed bars) and male rats, compared to wild-type rats (solid bars) and female rats, respectively. All animals were then then tested on spontaneous alternation (SA), social recognition (SR), and delayed non-matching-to-position (DNMTP) at 3-, 6-, and 9-months old. After behavioral testing at 9-months was completed, all rats were tested in activity chambers in order to measure locomotor activity (∼10 months old). All rats were euthanized, and tissue was processed for protein levels in the hippocampus.

### 2.3 Behavioral Assessment at 3-, 6-, and 9-Months Old

Utilizing a longitudinal design, all rats were tested on all three behavioral tasks at each age time point (3, 6, and 9 months; see *Figure 1* for timeline). Spontaneous alternation was utilized as an assay for spatial working memory, as deficits in this type of task have been found in the AD-rat model beginning at 6 months of age. Social recognition was used as a measure of ventral hippocampus-dependent social memory due to a previous study finding impairments in the AD-rats at 3 months of age, but not at other ages (Chaney et al., 2021). A delayed non-matching-to-position task was used as a previous study in AD-rats suggested a deficit in learning task rules, but not in short term delay-dependent memory, at 5-months of age (Muñoz-Moreno et al., 2018). Due to the previously reported anxiety of the F344 strain and low motor activity in the AD-rat model (Bert et al., 2002; Rex et al., 1996; Voorhees et al., 2019; Wu et al., 2020), activity chambers were utilized to measure general locomotion as well as investigative behaviors in an open arena at 10 months of age.

#### 2.3.1 Spontaneous Alternation

All rats were food restricted to 85-90% of their determined free-feeding weight and handled for seven days prior to testing. Testing for spontaneous alternation was performed using an elevated plus maze (105.5cm x 14.4cm x 15cm) with clear exterior walls. The testing room had clearly distinct spatial cues adhered to the walls as to guide the rats through the maze. All spatial cues remained constant for each rat. Rats were isolated and allowed to habituate to the room for 30 minutes before prior to testing. During testing, the rat was placed into the center of the plus maze and allowed to freely explore for 18 minutes. During this time, unique arm entries (defined as all four paws within an arm) were recorded. One complete spontaneous alternation is defined as four consecutively unique arm entries. Performance was recorded as a percentage of correct alternations, or the ratio of actual alternations to possible alternations (total alternations/[total arm entries-3] x100). After testing was complete, the rats were returned to their home cage and given free access to food to return to their free-feeding weights.

#### 2.3.2 Social Recognition

At 48 hours following the completion of spontaneous alternation, rats underwent social recognition testing (protocol modified from Mathiasen & DiCamillo, 2010). Testing for social recognition was performed in a clear plastic arena (58.4cm x 41.3cm x 31.4cm) with standard woodchip bedding. Rats (one adult and two juveniles) were acclimated to the testing room separately in clean standard housing cages (48.3cm x 26.7cm x 20.3cm) for 30 minutes. Adults and juveniles were kept separate to avoid olfactory contact. The two juveniles (50-80g; ∼1 month old F344 rats of same sex as adult experimental rat) used for recognition testing were not cage-mates and were marked for identification with different colors of low-odor semi-permanent animal markers (Stoelting, Wood Dale, IL, USA). After the initial acclimation to the room, the adult rat was placed into test arena 1 for an additional 30-minute acclimation to the arena.

A video camera placed above the area recorded the encoding trial (trial 1). Juvenile 1 was introduced into the test area 1, with the adult rat allowed to freely explore the juvenile for 5 minutes. At the end of trial 1, the juvenile was removed from test arena 1 and placed back into its acclimation cage. The adult was then transferred to a new arena with fresh bedding (test arena 2) for a 30-minute inter-trial interval. The retention trial (trial 2) then took place and was recorded. Juvenile 1 (familiar) and juvenile 2 (novel) were placed into test arena 2 with the adult rat for 5 minutes. After trial 2, testing was complete, and all rats were placed back into their home cages.

Video recordings of each trial were analyzed by blind experimenters. Experimenters followed a detailed protocol and were extensively trained to ensure strong inter-rater reliability. A stopwatch was used to determine the total time spent interacting with each juvenile. Social interaction was defined as the adult grooming, sniffing, pawing, or having any close contact (nose, mouth, paws) with the juvenile. An exploration ratio was determined for each adult by dividing the time spent investigating juvenile 2 (novel) by the total time investigating both juvenile 1 and 2 during trial 2. An exploration ratio of 0.5 indicates equal time spent with each juvenile, suggesting a lack of recognition of the familiar juvenile 1. An exploration ratio greater than 0.5 indicates more time spent with the novel juvenile, suggesting some recognition of the familiar juvenile 1.

#### 2.3.3 Delayed Non-Matching-to-Position (DNMTP)

Following the completion of social recognition testing, all rats were placed back on food restriction and gradually restricted to 85% of their free-feeding weight for operant training.

Weights were maintained at 85-90% of their baseline throughout operant testing while ensuring they never reached levels lower than 85% of the standard growth curve of the F344 strain. Rats were exposed to the operant food reward (Rodent Purified Dustless Precision Pellet; Bio-Serve, Flemington, NJ, USA) in their home cage for two consecutive days before operant training began. Operant chambers (30 cm × 33 cm × 23 cm; Med Associates Inc., St. Albans, VT, USA) were enclosed within sound attenuating boxes (59 cm × 55 cm × 36 cm) with constantly running fans for white noise. All operant chambers were equipped with two retractable levers with a stimulus light above each. A magazine for dispensing the food reward was placed evenly between the two levers. A house light for general illumination of the box was located on the opposite wall of the levers and magazine. Operant boxes were interfaced with MED-PC software (Med Associates Inc. St Albans, VT, USA).

The DNMTP task consisted of four phases: habituation and magazine training, pretraining for lever pressing, zero-delay NMTP, and finally delayed NMTP. All rats were habituated to the operant box for 30 minutes, followed by 30 minutes of magazine training where the rat learned to retrieve the food reward from the magazine. During the pretraining phase, rats were behaviorally shaped to lever press on a fixed ratio 1 (FR1) schedule of reinforcement. Each rat was randomly assigned to initially learn to press either the left or right lever. During this pretraining exercise, the lever did not retract and can be pressed continuously for a food reward. Once the criterion was met (50 reinforced lever presses during a single session), the rat moved on to learn to lever press the opposite lever under the same criteria. Once shaping was complete, rats were trained to press the lever within a 10-second period between extension and retraction of the lever. Extension of the left or right lever was randomized. If the rat failed to press the lever in that 10-second time period, it was counted as an omission. Rats needed to have fewer than five omissions for two consecutive days before beginning 0-delay NMTP training.

##### Zero Delay NMTP

During a sample phase, either the left or right lever was extended for the rat to press for a food reward. The presentation of the levers was pseudorandomized, and the rat had 1 second to respond before retraction. If the rat did not press the lever during a trial, a 30-second intertrial interval occurred in which the house light was shut off. If the rat pressed the extended lever, the lever retracted, and a food reward was given. Once a nose poke occurred in the magazine, both levers were extended for a choice response phase. The rat had to choose the opposite (non-matching) lever from what was extended in the sample phase in order to pass the trial and obtain another food reward. Each session included 72 trials and the criterion was met when a rat achieved 90% or higher for two consecutive sessions. During retraining at 6- and 9- months of age, rats only needed to achieve 90% or higher for one session to move on to delayed NMTP.

##### Delayed NMTP

Once a rat met the criteria for learning 0-delay NMTP, they were tested on delayed NMTP (DNMTP) for 10 consecutive days, regardless of performance. The same ITI and omission parameters as zero delay NMTP were followed for DNMTP. However, when the nose poke into the magazine occurred following the sample phase, a randomly selected delay of 0, 4, 8, or 16 seconds occurred before the subsequent extension of both levers for the choice response phase. Each delay interval was randomly distributed across 72 trials within a single session. Total percentage of correct NMTP choices for each delay timepoint was determined and recorded daily for each rat. Once complete, rats were returned to their home cages with free access to food. Rats remained in standard housing conditions until the next round of behavioral testing.

#### 2.3.4 Activity Chambers

Following completion of the 9-month DNMTP testing (∼10 months old), rats were returned to their home cages and given free access to food. Once their weight recovered, activity chambers (Accuscan Instruments, Columbus, OH, USA) were used to assess any changes in generalized locomotor activity due to genotype or ethanol exposure. Six acrylic chambers (41 x 41 x 30.5cm) were each surrounded by a 15x15 infrared photocell arrays and were synched with Versamax and Versadat software (Accuscan Instruments) to measure and record patterns of horizontal and vertical beam breaks. The total time spent moving (in seconds), time spent in the margins of the chamber (in seconds), and the time spent in the center of the chamber (in seconds) were recorded every 5 minutes for a period of 30 minutes. All testing occurred between 1:00 PM and 3:00 PM.

### 2.4 Tissue Preparation

After completion of all behavioral tasks (∼11 months of age), rats were euthanized by rapid decapitation. At the time of decapitation and dissection, brains from female rats did not differ in weight between genotypes or exposure conditions (*p*’s > 0.24). For male rats, brains from AD-AIE exposed rats (*M* = 1.956 grams, *SD* = 0.046) weighed less than brains from WT control rats (*M* = 2.004 grams, *SD* = 0.058; *p* = 0.038). Whole brains were flash frozen by a 2-minute submersion in -20°C methyl butane (temperature maintained by dry ice) and then stored at -80°C until tissue punching. The MS/DB and hippocampus were bilaterally micropunched (1.0–2.0 mm; EMS-Core Sampling Tools, Electron Microscopy Sciences, Hatfield, PA, USA) using a cryostat collection procedure with the temperature maintained at -20°C. Coordinate ranges for the MS/DB and hippocampus tissue punches were determined using a rat brain atlas (Paxinos and Watson, 2013) and pilot brain tissue and were consistent across all rats.

### 2.5 Western Blot Analysis in the Dorsal and Ventral Hippocampus

The hippocampus of each animal was homogenized in a lysis buffer (1% SDS, 1 mM EDTA, 10 mM Tris) containing protease inhibitors (Halt™ Protease Inhibitor Cocktail, ThermoFisher Scientific, Waltham, MA, USA) and phosphatase inhibitor cocktails 2 and 3 (Sigma-Aldrich, St. Louis, MO, USA) to prevent degradation of the protein of interest. Protein concentrations were determined using a bicinchoninic acid method (Pierce, Rockford, IL, USA) and compared to standards of bovine serum albumin. Samples were normalized and, as determined by linear range test (data not shown) 10–20 μg of total protein from dorsal hippocampus and 20-30 μg of total protein from ventral hippocampus were denatured and separated using electrophoresis on Novex™ 8–16% Tris-Glycine sodium dodecyl sulfate polyacrylamide gels (Invitrogen, Carlsbad, CA, USA), and were subsequently transferred to a polyvinylidene difluoride membrane (Invitrogen, Carlsbad, CA, USA). Each membrane underwent a Total Protein Stain Kit (TotalStain Q, Azure Biosystems, Dublin, CA, USA) and was blocked for one hour in 5% BSA, 0.01% Tween-20 in TBS. The first set of membranes were first probed with an antibody for pTau (Thr181), mildly stripped with Restore stripping buffer (ThermoFisher Scientific), and then re-probed for total Tau (Tau46), stripped again and re-probed with B-Actin. To look at all of the proteins of interest and avoid cross-reactivity, the blots were cut into four strips and each strip was incubated with primary antibodies for TrkA, vAChT, B-actin, and mature NGF (mNGF) and its precursor, proNGF. Another set of membrane strips were incubated with primary antibodies for tropomyosin-related kinase B receptor (TrkB), p75NTR, B-Actin, or mature Brain-derived neurotrophic factor (BDNF) and its precursor, proBDNF. The following day, blots were incubated in a peroxidase-conjugated secondary antibody (see Table 1 for all antibodies) for one hour. Protein levels were detected with enhanced chemiluminescence (Pierce™ ECL Western Blotting Substrate, ThermoFisher Scientific). Β-actin and total protein were analyzed, but total protein was used for normalization in all analyses. Levels of pTau were analyzed as a measure of pTau/total Tau/housekeeper (total protein). A cross-reactivity test with primary antibodies for pTau, total Tau, and β-actin and their secondary antibodies was carried out to confirm that there was no false detection (data not shown). All gels were run in duplicate with the optical band densities being averaged. All gels were separate by sex and counterbalanced across exposure and genotype conditions.

**Table 1.** Primary and Secondary Antibodies for Western Blots.

| **Protein** | **Primary Antibody** | **Secondary Antibody** |
| --- | --- | --- |
| NGF; proNGF | Rabbit 1:200 (Abcam; ab52618) | Goat 1:250 (Abcam; ab205718) |
| BDNF, proBDNF | Rabbit 1:200 (Abcam; ab108319) | Goat 1:250 (Abcam; ab205718) |
| TrkA | Rabbit 1:1000 (Abcam; ab302524) | Goat 1:1000 (Abcam; ab205718) |
| TrkB | Rabbit 1:1000 (Abcam; 187041) | Goat 1:2000 (Abcam; ab205718) |
| VACHT | Rabbit 1:1000 (Synaptic Systems; 139103) | Goat 1:1000 (Abcam; ab205718) |
| p75NTR | Rabbit 1:300 (EMD; 07-476) | Goat 1:500 (Abcam; ab205718) |
| pTau | Rabbit mAb 1:1000 (CST; 12885) | Goat 1:1000 (Abcam; ab205718) |
| Tau 46 | Mouse mAb 1:1000 (CST; 4019) | Goat 1:10000 (Invitrogen; A16078) |
| β-actin | Chicken 1:2000 (Origene; TA349013) | Donkey 1:1000 (Invitrogen; SA1-300) |

*Note.* Host, dilution, company, and catalog information. Nerve growth factor (NGF, pro-NGF); brain-derived neurotrophic factor (BDNF; proBDNF); vesicular acetylcholine transporter (VACHT); tropomyosin receptor kinase A (TrkA) and B (TrkB); p75 neurotrophic receptor (p75NTR); Beta-actin ( β-actin); phospho-tau [Thr181; D9F4G] (pTau).

### 2.6 Statistical Analyses

Analyses were performed in GraphPad Prism (version 9.5.1). All analyses were separated by sex, due to initial differences in blood ethanol concentrations and to reduce any spurious effects from a 4- or 5-way ANOVA. A repeated measure factorial ANOVA (Exposure [AIE, CON] x Genotype [AD, WT] x Age [3-, 6-, 9-Months; within subjects]) was used to assess spontaneous alternation scores and arm entries in female rats. Male rats at 9 months of age did not meet criterion for entering at least 10 arms during spontaneous alternation; therefore, a repeated measures factorial ANOVA (Exposure [AIE, CON] x Genotype [AD, WT] x Age [3-, 6-Months; within subjects]) was used to assess spontaneous alternation scores and arm entries in male rats. A repeated measures factorial ANOVA (separated by Sex; Exposure [AIE, CON] x Genotype [AD, WT] x Age [3-, 6-, 9-Months; within subjects]) was used to assess social recognition discrimination ratios. *T*-tests were used to determine differences in social recognition discrimination ratios from a theoretical value of 0.50 (equal time spent with each juvenile rat).

Data from the DNMTP task were separated by sex, age, and test day in order to analyze the results in a meaningful way without the heightened risk of spurious effects. Days 1, 5, and 10 were analyzed as measures of beginning, middle, and ending performance across all 10 test days. Individual repeated measure ANOVAS (separated by sex, age, and day; Exposure [AIE, CON] x Genotype [AD, WT], Delay [0-, 4-, 8-, 16-seconds; within subjects]) were used to examine percent of correct choices on the DNMTP task. The number of zero delay NMTP trials to meet criterion was analyzed using a univariate ANOVA (separated by day and sex; Exposure [AIE, CON] x Genotype [AD, WT]). Separate univariate ANOVAs (separated by sex; Exposure [AIE, CON] x Genotype [AD, WT]) were used to examine activity data (movement time, total distance and center margin ratio separated) and protein levels in each brain region of interest. When appropriate, Fisher’s LSD test was used for post-hoc analyses. Data points two standard deviations above or below the initial group mean were removed as outliers.

## 3 Results

### 3.1 Blood Ethanol Concentrations

Blood sample analyses show that rats exposed to AIE reached heavy to extreme binge-like levels of intoxication (greater than 80mg/dL; Spear, 2018). AD rats (*M* = 229.70, *SD* = 19.14) had higher BECs than WT rats (*M* = 194.90, *SD* = 10.04; *F*[1,54] = 11.33, *p* = 0.001; see *Figure 1*). Male rats (*M* = 222.61, *SD* = 29.09) had higher BECs than female rats (*M* = 201.97, *SD* = 20.00; *F*[1,54] = 4.00, *p* = 0.05).

### 3.2 Spontaneous Alternation

#### Female data

Data points from six female rats were excluded due to failure to meet task demands at all age time-points. Final number of female rats per condition were: WT (10 CON, 10 AIE), AD (7 CON, 7 AIE). There was an Age X Exposure interaction on percent alternation in female rats (*F*[2,60] = 4.36, *p* = 0.017; see *Figure 2A*). At 3-months of age, all female groups performed comparably on the spontaneous alternation task. However, by 6-months of age, selectively TgF344-AD AIE-exposed female rats were impaired compared to both water control (*p* = 0.008) and AIE-WT rats (*p* = 0.001). At 9-months of age, we see that AIE leads to lower percent alternation scores in both TgF344-AD (*p* = 0.016) and wildtype (*p* = 0.006) AIE-exposed rats, relative to WT controls.

**Figure 2.**
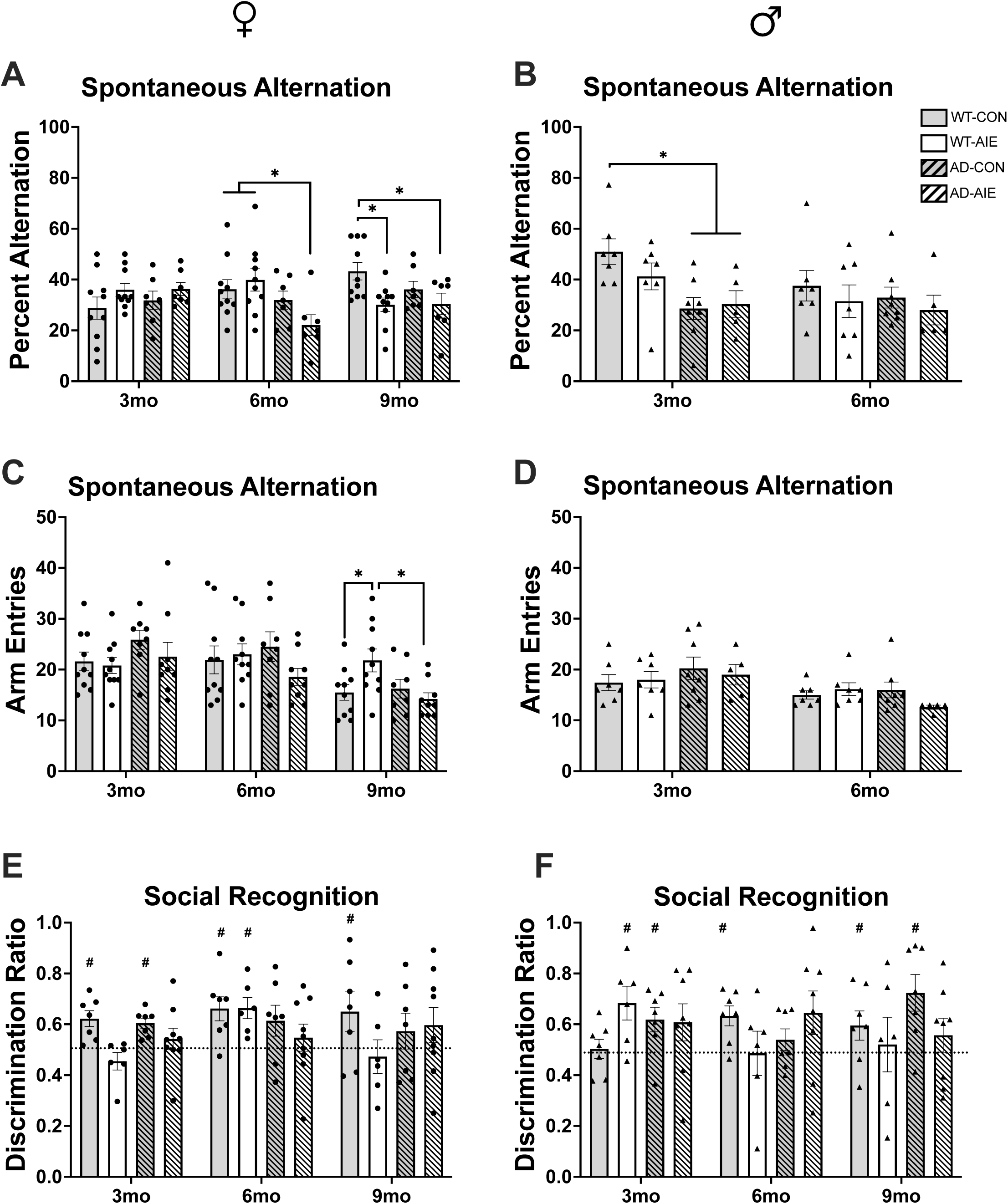
Spontaneous Alternation and Social Recognition data. Spontaneous alternation scores for female rats (A) and male rats (B). Arm entries on the spontaneous alternation task for female rats (C) and male rats (D). Circle data points represent female rats; triangle data points represent male rats. Solid gray bars represent wild-type water control rats; solid white represent wild-type AIE-exposed rats, dashed gray bars represent water control TgF344-AD rats; dashed white bars represent AIE-exposed TgF344-AD rats. * = *p* < 0.05; ** = *p* < 0.01 Social recognition scores for female rats (E) and male rats (F). Circle data points represent female rats; triangle data points represent male rats. # = significant difference from a theoretical “chance” value of 0.50.

Female rats also displayed an effect of Age (*F*[1,33] = 12.66, *p* < 0.0001) on total arm entries (see *Figure 2C*). There was also an Age X Genotype interaction (*F*[2,66] = 3.37, *p* =0.04) on arm entries, with every group except for WT-AIE female rats making fewer arm entries as they age.

#### Male data

Data points from 13 male rats were excluded due to failure to meet task demands at all age time-points. Final number of male rats per condition were: wild-type (7 water control, 7 AIE), TgF344-AD (8 water control, 5 AIE). Male AD-rats performed worse than WT male rats on spontaneous alternation (effect of Genotype; *F*[1,23] = 6.49, *p* = 0.018; see *Figure 2B*). There was no effect of Age or Exposure on alternation scores (both *p*’s > 0.19).

However, arm entries on the spontaneous alternation task declined with age in male rats (effect of Age; (*F*[1,23] = 10.03, *p* = 0.004; see *Figure 2D*). By 6 months of Age 24%, of rats did not reach the criterion of 10 arm entries, and by and this increase to 56% by 9 months. There was no effect of Genotype or Exposure on arm entries in male rats.

### 3.3 Social Recognition

Data points from seven female rats were excluded due to failure to meet task demands at all age time-points. Final number of female rats per condition were: WT (7 CON, 6 AIE), AD (7 CON, 9 AIE). Eight male rats were excluded due to failure to meet task demands at all age time-points and the final number of male rats per condition were: WT (7 CON, 6 AIE), AD (7 CON, 8 AIE).

### Group differences

#### Female Data

There was a significant Exposure effect on social recognition discrimination ratios (DR) in female rats (*F*[1,25] = 4.68, *p* = 0.04), with AIE-exposed rats performing worse than CON rats overall. There were no effects of Age or Genotype on social recognition discrimination ratios in female rats (*p*’s > 0.21).

#### Male Data

There were no effects of Exposure, Genotype, or Age on social recognition discrimination ratios in male rats (*p*’s > 0.22).

### Difference from chance performance

Discrimination ratios (DR) above 0.5 (as determined by *t*-tests) indicate more time spent with the novel juvenile, which is interpreted as intact memory for the familiar juvenile. When DRs are not significantly above 0.5, this implies equal time with the novel and the familiar juvenile. This is interpreted as impaired social memory, or also considered discriminating novel conspecifics at “chance level”.

#### Female Data

At 3-months of age, AIE-exposed, both WT and AD, female rats performed at chance level on social recognition, while both WT (*p* = 0.001) and AD (*p* = 0.004) CON rats showed a preference for the novel juvenile (see *Figure 2E).* At 6-months of age, WT female rats showed a preference for the novel juvenile, while AD female rats perform at chance level. At 9-months of age, only the WT CON female rats spent significantly more time with the novel juvenile over the familiar juvenile (*p* = 0.048); all other groups spent equal amounts of time with each juvenile, demonstrating loss of social recognition memory.

#### Male Data

Interestingly, only WT-AIE (*p* = 0.049) and AD-CON (*p* = 0.014) male rats performed significantly above chance on social recognition at 3-months old (*Figure 2F)*. At 6-months old, only the WT-CON rats performed above chance (*p* = 0.034). Finally, at 9-months old, only WT-CON (*p* = 0.049) and AD-CON male rats performed above chance (*p* = 0.012).

### 3.4 Delayed Nonmatching to Position (DNMTP) Zero delay acquisition—Trials to criterion

#### Female Data

There were no effects of Exposure or Genotype on zero delay NMTP trials to criterion in female rats at 3-months old (WT-CON=855(mean)±156(SEM); WT-AIE=792±92; AD-CON=639±77; AD-AIE=675±80; all *p*’s > 0.12). At 6-months of age, AD female rats took less trials to re-establish criterion than WT rats (effect of Genotype: *F*[1,40] = 7.65, *p* = 0.008; WT-CON=176±32; WT-AIE=168±15; AD-CON=111±77; AD-AIE=114±80). Exposure to ethanol did not affect trials to criterion at 6-months old in female rats (*p* = 0.90). There were no effects of Exposure or Genotype to re-establish criterion in female rats at 9-months of age (all *p*’s > 0.42; WT-CON=80±8; WT-AIE=96±10; AD-CON=98±15; AD-AIE=96±10).

#### Male Data

See *Figure 5* for individual analyses at 3-months old (A), 6-months old (B), and 9-months old (C). At 3-months of age, there was a nonsignificant trend towards an effect of Genotype, with AD male rats taking slightly more trials to meet criterion than WT rats (*F*[1,28] = 3.27; *p* = 0.072; WT-CON=450±23; WT-AIE=369±25; AD-CON=567±99; AD-AIE=459±36).

There were no effects of Exposure or Genotype in male rats at 6-months old to re-establish criterion (all *p*’s > 0.39; WT-CON=104±15; WT-AIE=104±11; AD-CON=98±11; AD-AIE=120±10). At 9-months of age, AIE-exposed male rats took more trials to re-establish criterion than CON male rats (effect of Exposure: *F*[1,38] = 5.58, *p* = 0.023; WT-CON=72±5; WT-AIE=85±9; AD-CON=72±3; AD-AIE=101±16). Genotype did not affect trials to criterion at 9-months old in male rats (*p* = 0.38).

### Delayed NMTP

All groups showed an effect of delay (all *p*’s < 0.001): as the delay interval increases, the percentage of correct lever choices decreases. There were no clear patterns of AIE exposure or genotype affecting DNMTP position in either sex across days or ages (see *Figures 3* and *4*).

**Figure 3.**
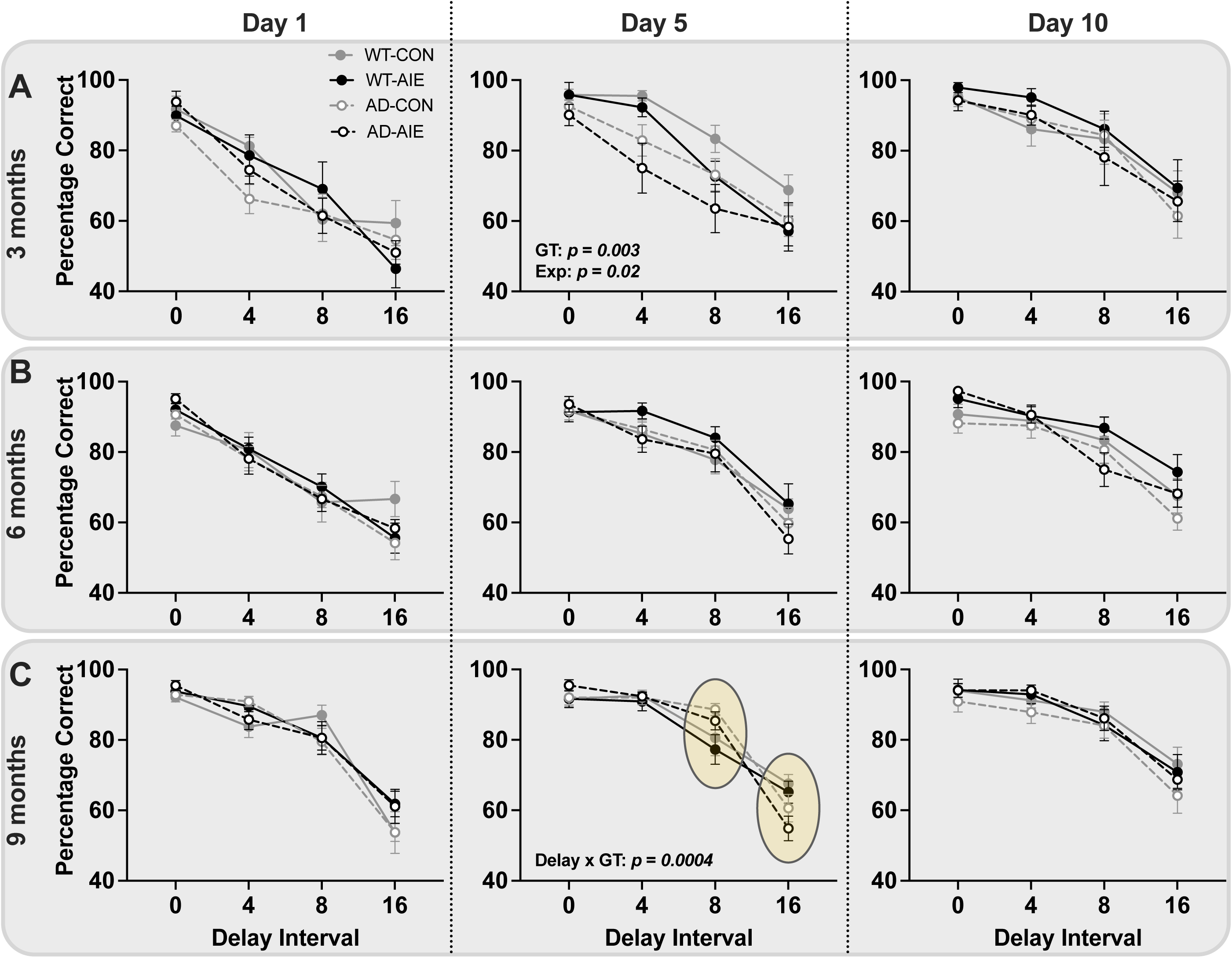
**Delayed non-matching to position performance for female rats** at 3-months old (A), 6-months old (B), and 9-months old (C). Solid gray lines represent wild-type water control female rats; solid black lines represent wild-type AIE-exposed female rats, dashed gray lines represent water control TgF344-AD female rats; dashed black lines represent AIE-exposed TgF344-AD female rats. Highlighted regions emphasize interactions that are present in the data (listed on graph).

**Figure 4.**
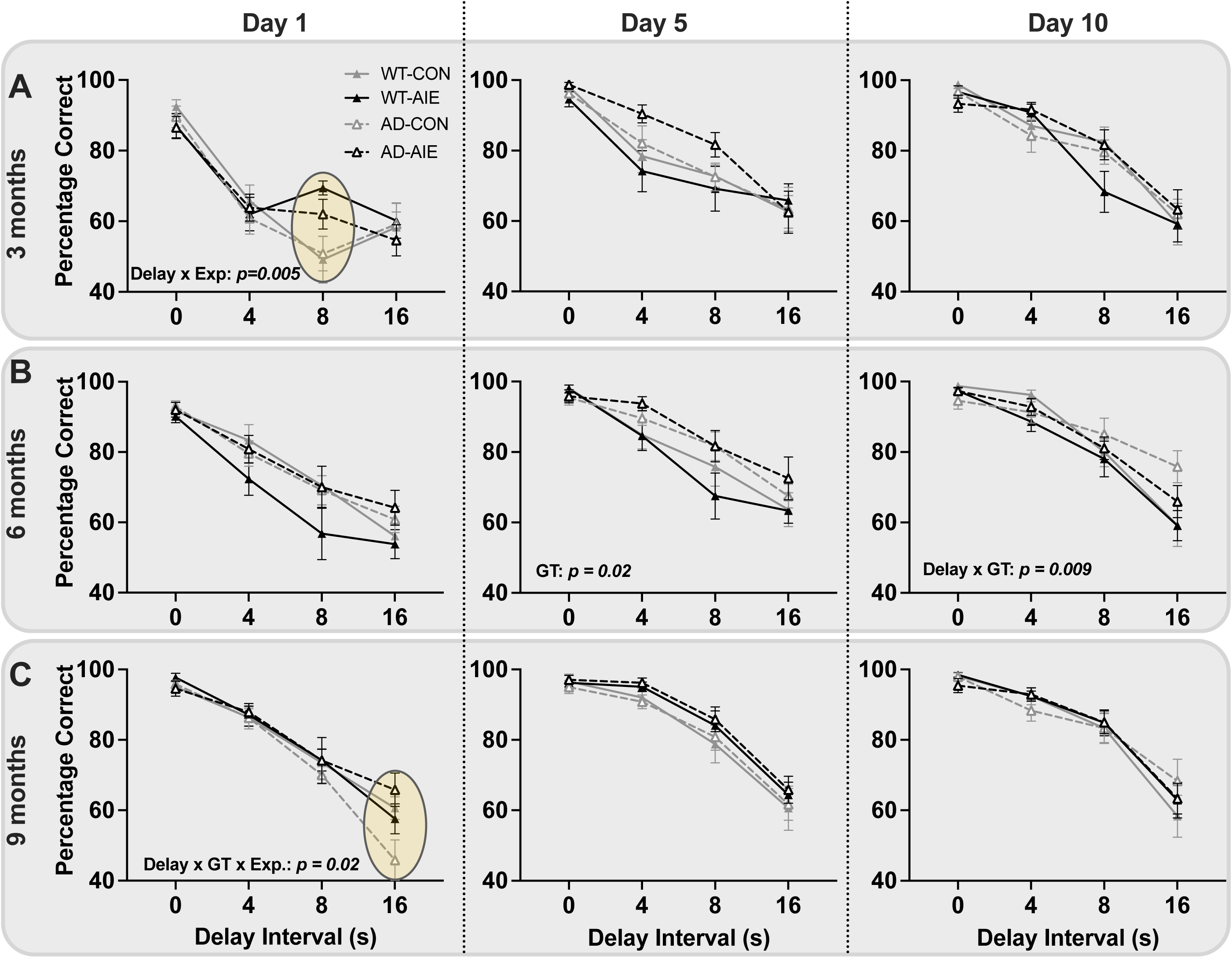
**Delayed non-matching to position results for male rats** at 3-months old (A), 6-months old (B), and 9-months old (C). Solid gray lines represent wild-type water control male rats; solid black lines represent wild-type AIE-exposed male rats, dashed gray lines represent water control TgF344-AD male rats; dashed black lines represent AIE-exposed TgF344-AD male rats. Each session of testing (1/day) was 72 trials. Highlighted regions emphasize interactions that are present in the data (listed on graph).

#### Female Data

See *Figure 3* for individual analyses at 3-months old (A), 6-months old (B), and 9-months old (C). There was an effect of Genotype (*F*[1,28] = 10.45, *p* = 0.003) and Exposure (*F*[1,28] = 5.47, *p* = 0.02) on Day 5 in 3-month-old female rats: CON rats performed better than AIE-exposed rats and WT rats performed better than AD rats. There was a Delay X Genotype interaction on Day 5 in 9-month-old rats (*F*[3,117] = 6.54, *p* = 0.0004).

#### Male Data

See *Figure 4* for individual analyses at 3-months old (A), 6-months old (B), and 9-months old (C). There was a Delay X Exposure interaction on Day 1 of 3-month-old rats (*F*[3,102] = 4.45, *p* = 0.005), with AIE-exposed rats performing better at the 8-second delay than CON rats. There was a Genotype effect on Day 5 in 6-month-old rats, with AD rats performing better than WT rats (*F*[1,37] = 5.35, *p* = 0.02). There was a Delay X Genotype effect on Day 10 in 6-month-old rats (*F*[3,114] = 4.00, *p* = 0.009). There was a Delay X Genotype X Exposure interaction on Day 1 in 9-month-old rats (*F*[3,114] = 3.26, *p* = 0.02). Finally, there was a trend towards an effect of Exposure on Day 5 in 9-month-old rats, with AIE-exposed rats performing slightly better than CON rats (*p* = 0.06).

In summary, short-term memory performance was decreased by AIE and AD transgenes in female rats. In contrast, the opposite effect was found in male rats: AIE and AD transgenes improved short-term memory performance.

### 3.5 Activity Chambers

All rats were assessed at 11 months of age for locomotor ability using standard activity chambers. Several key activity measures were recorded and analyzed to determine if AD transgenes or adolescent ethanol exposure altered general activity levels.

*Movement Time (s):* Female AD rats moved for less time than wildtype female rats (effect of Genotype; *F*[1,38] = 4.40, *p* = 0.043; see *Figure 5A*), specifically in the CON condition (*p* = 0.047). There was no effect of Exposure or Genotype on movement time in male rats.

**Figure 5.**
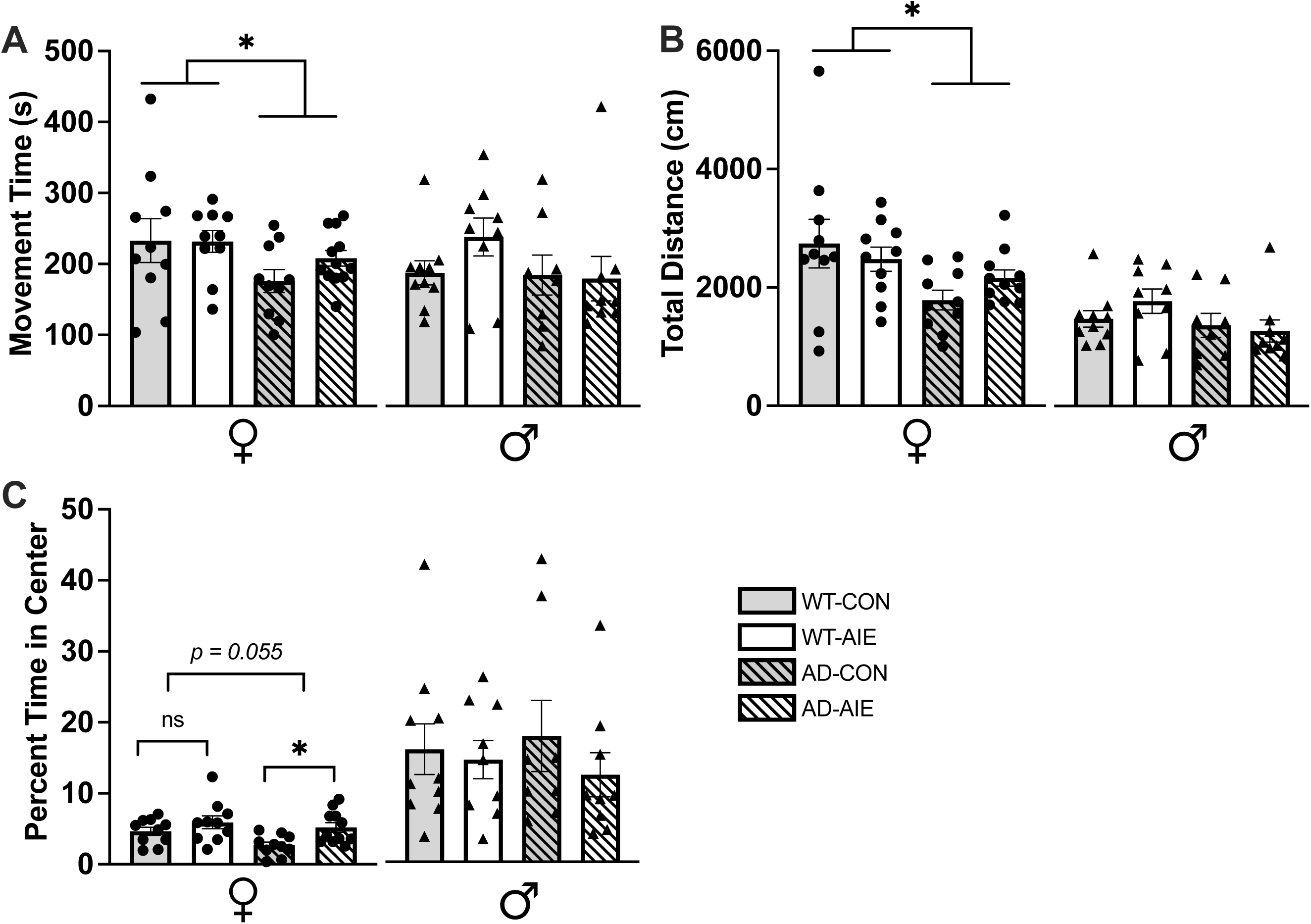
Analyses of locomotor activity from activity chambers at 10-months of age. A) total movement time (s; out of 1800 s), B) total distance moved (cm), and C) percentage of time in the center of the arena. Circle data points represent female rats; triangle data points represent male rats. Solid gray bars represent wild-type water control rats; solid white bars represent wild-type AIE-exposed rats, dashed gray bars represent water control TgF344-AD rats; dashed white bars represent AIE-exposed TgF344-AD rats. * = *p* < 0.05

*Total Distance (cm):* Female AD-rats had a lower total distance moved than WT female rats (effect of Genotype; *F*[1,38] = 6.57, *p* = 0.015; see *Figure 5B*), specifically in the CON condition (*p* = 0.01). There was no effect of Exposure or Genotype on total distance moved in male rats.

*Time in Center:* A percentage of time spent in the center of the activity chamber was calculated by dividing time spent in the center by total time in the chamber (1800 seconds). There was an effect of Exposure effect in female rats (*F*[1,37] = 7.63, *p* = 0.009; see *Figure 5C*), driven by AD-CON rats spending less time in the center of the chamber than AD-AIE exposed female rats (*p* = 0.012). There was a trend towards a significant Genotype effect, with AD female rats spending slightly less time in the center of the chamber than WT female rats (*F*[1,37] = 3.90, *p* = 0.055). There was no effect of Exposure or Genotype on center-to-margin ratios in male rats.

### 3.6 Protein Levels in the Dorsal and Ventral Hippocampus

The differences in protein levels are expressed as percent of WT control separately for female and male WT and AD rats as a function of Genotype and Exposure.

*pTau (Thr181):* There was no effect of Genotype or Exposure on levels of pTau in the dorsal or ventral hippocampus of female rats (all *p*’s > 0.16; see *Figures 6A&C*). However, in male rats there was a main effect of Exposure on levels of pTau in the dorsal hippocampus with AIE-exposed male rats having higher levels than CON male rats (*F*[1,33] = 6.88, *p* = 0.013; see *Figures 6B&D*). In contrast, in the ventral hippocampus, not the dorsal hippocampus (*p*=0.67), there was an effect of Genotype on pTau expression in male rats (*p* = 0.03), with male rats with AD transgeneses having higher levels of pTau.

**Figure 6.**
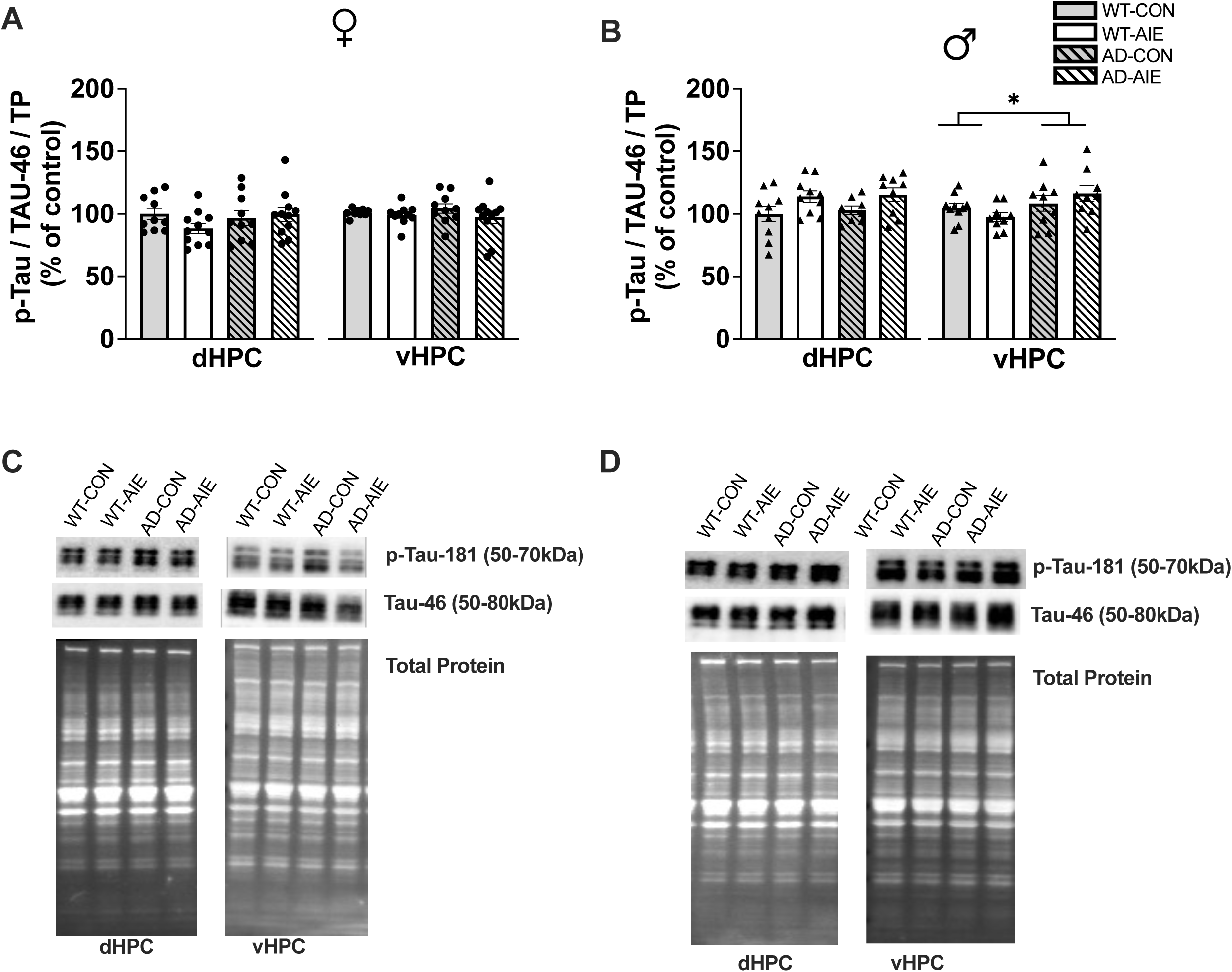
**pTau (Thr181) protein levels in the dorsal (dHPC) and ventral hippocampus (vHPC)** of female rats (AC) and male rats (BD). In the ventral hippocampus, male AD rats had higher levels of pTau. *GT= main effect of Genotype; *Exp=main effect of Exposure; *GT x Exp= Interaction of Genotype x Exposure

*TrkA:* Female rats exposed to AIE had a lower expression of TrkA in the dorsal hippocampus than CON female rats (effect of Exposure; *F*[1,31] = 8.63, *p* = 0.006), an effect not seen in the ventral hippocampus (*p*’s > 0.17; see *Figure 7A&I*). Overall, male rats exposed to AIE had a lower expression of TrkA than CON male rats across the dorsal and ventral hippocampus (effect of Exposure; *p*= 0.01). Although there was no effect of Genotype on TrkA expression in the dorsal hippocampus in male rats (*p* = 0.69), in the ventral hippocampus male AD rats had higher levels of TrkA expression than WT rats (*p* = 0.036; see *Figure 7B&J*).

*VAChT*: AD-female rats had lower levels of VAChT than WT female rats in the dorsal hippocampus (effect of Genotype; *F*[1,33] = 21.18, *p* < 0.0001), that was not seen in the ventral hippocampus (*p*=0.86). Exposure to AIE did not affect VAChT levels in the dorsal or ventral hippocampus of female rats (all *p’s* > 0.08; see *Figure 7C&I*).

One data point (wild-type AIE) was removed as an outlier for male rats. There was an interaction between Genotype and Exposure on levels of VAChT in the dorsal hippocampus (*F*[1,30] = 5.14, *p* = 0.031). In AD male rats, AIE-exposed animals had lower levels of vAChT than AD-CON male rats. There were no effects of interactions between Exposure and Genotype in the ventral hippocampus (all *p’s* > 0.26; see *Figure 7D&J*).

*proNGF and mNGF:* Protein levels for mature (m) and proNGF were not analyzed in the dorsal hippocampus of female rats due to initial technical difficulties and low sample volumes. In the ventral hippocampus, there were no differences in proNGF levels has a function of Exposure or Genotype (all *p*’s > 0.73) in female rats (see *Figure 7E&I*). However, in the ventral hippocampus of female rats there was an interaction between Exposure and Genotype (*F*[1,37] = 5.43; *p* = 0.025) for mNGF levels: Female AD-AIE rats had lower levels of mNGF than AD-CON rats (*p =* 0.036; see *Figure 7G&I*).

In the dorsal hippocampus, male rats exposed to AIE had a lower expression of proNGF than CON rats (effect of Exposure; *F*[1,31] = 8.23, *p* = 0.007). There was a trend towards a Genotype effect with AD male rats having higher levels of proNGF than WT male rats (*F*[1,31] = 3.87, *p* = 0.058). In the ventral hippocampus, there was an interaction between Exposure and Genotype on proNGF levels (*F*[1,34] = 7.40; *p* = 0.01): Male AD-AIE rats had the highest level of proNGF(*p* = 0.002; see *Figure 7F&J*). In the dorsal hippocampus, there was no effect of Genotype or Exposure on levels of mNGF in the hippocampus of male rats (*p*’s > 0.42).

However, in the ventral hippocampus, male rats with AD transgenes had lower levels of mNGF (effect of Genotype; *F*[1,33] = 8.53; *p* = 0.006; see *Figure 7H&J*).

*TrkB:* Although there was no effect of Exposure or Genotype on TrkB expression in the dorsal hippocampus of female (all *p*’s > 0.38; see *Figure 88A&I)* or male rats (all *p*’s > 0.12; see *Figure 8B&J*), in the ventral hippocampus AD transgenes increased the level of TrkB expression in the ventral hippocampus of both male and female rats (both *F*[1,34] < 6.97; *p’s*<0.02).

**Figure 8.**
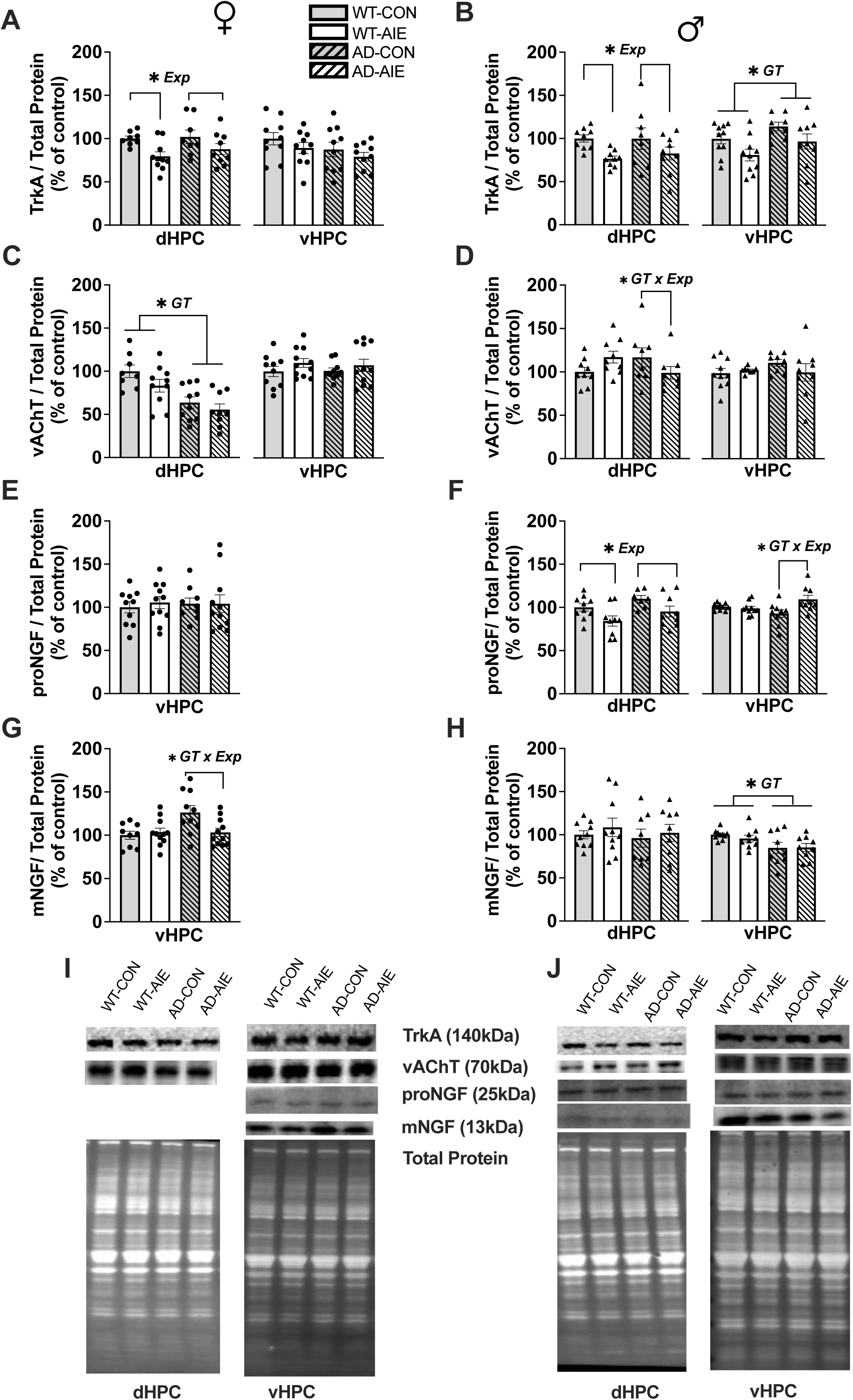
**Protein levels of (A) TrkA, (B) VAChT, (C) proNGF and (D) mNGF in the dorsal (dHPC) and ventral hippocampus (vHPC)**. Solid gray bars represent wild-type water control rats; solid white bars represent wild-type AIE-exposed rats, dashed gray bars represent water control TgF344-AD rats; dashed white bars represent AIE-exposed TgF344-AD rats. *GT= main effect of Genotype; *Exp=main effect of Exposure; *GT x Exp= Interaction of Genotype x Exposure

**Figure 8.**
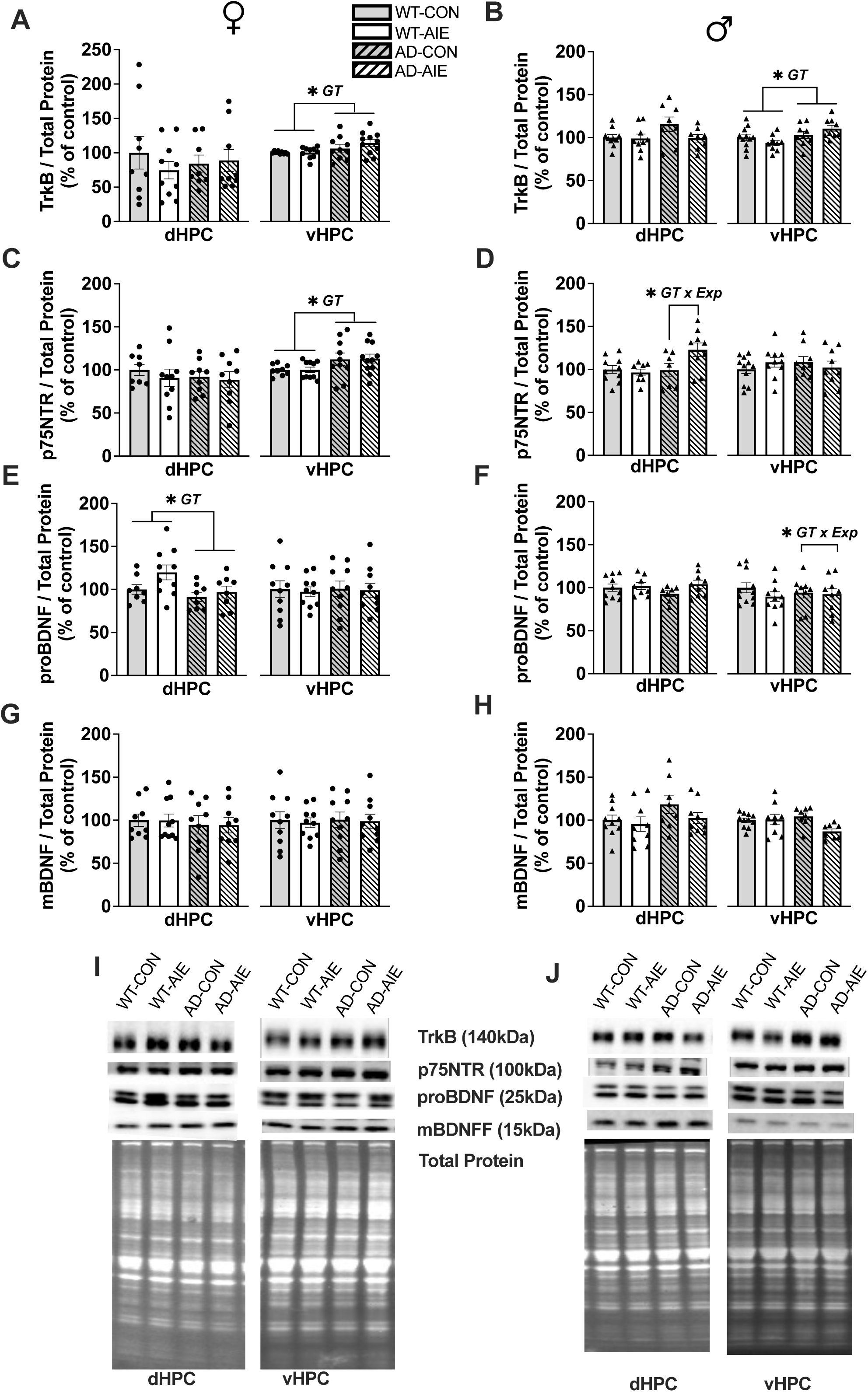
Protein levels in the hippocampus of male rats. A) VAChT protein levels, B) p75 protein levels, C) pTau-181 protein levels, D) proBDNF protein levels, E) BDNF protein levels, F) TrkB protein levels, G) TrkA protein levels, H) proNGF protein levels, I) NGF protein levels, J) raw band images. Solid blue bars represent wild-type water control rats; solid red bars represent wild-type AIE-exposed rats, dashed blue bars represent water control TgF344-AD rats; dashed red bars represent AIE-exposed TgF344-AD rats. *GT= main effect of Genotype; *Exp=main effect of Exposure; *GT x Exp= Interaction of Genotype x Exposure

*p75NTR:* In the dorsal hippocampus there was no effect of Genotype or Exposure on levels of p75NTR in female rats (all *p*’s > 0.46). However, in the ventral hippocampus there was a significant effect of Genotype with AD female rats having higher p75NTR levels that WT female rats (*F*[1,36] = 6.49; *p* = 0.015; see *Figure 8C&I*). In male rats, there was an interaction between Genotype and Exposure (*F*[1,30] = 4.76, *p* = 0.037) in the dorsal hippocampus was driven by AD male rats exposed to AIE having higher levels of p75NTR than all other groups (*p* = 0.005). There were no differences as a function of Genotype or Exposure in the ventral hippocampus in male rats (all *p’s* > 0.22; see *Figure 8D&J*).

*proBDNF and mBDNF:* AD female rats had lower levels of proBDNF in the dorsal hippocampus compared to WT female rats (effect of Genotype; *F*[1,30] = 4.93, *p* = 0.034; see *Figure 8E&I*). There was no effect of Exposure or Genotype on levels of proBDNF in the ventral hippocampus of female rats. In addition, mBDNF levels across the hippocampus of female rats were unaffected by Exposure and Genotype (all *p*’s > 0.23; see *Figure 8G&I*).

In male rats, Exposure and Genotype did not influence proBDNF or mBDNF levels in the dorsal hippocampus (all *p*’s> 0.11). There was an interaction between Genotype and Exposure in the ventral hippocampus of male rats (*F*[1,32] = 5.06; *p* = 0.031) and there was a trend towards an effect of Exposure (*F*[1,32] = 4.08; *p* = 0.051), AD-AIE male rats had lower levels of proBDNF (*p*’s < 0.03; see *Figure 8F&J*) compared with other groups. However, across the hippocampus there were no differences as a function of Exposure and Genotype on mBDNF levels in male rats (all *p*’s > 0.11; see *Figure 8H&J*).

## 4 Discussion

A longitudinal design was used to characterize and map out behavioral impairments in a transgenic model of AD (TgF344-AD) following adolescent ethanol exposure. Alterations in AD-related gene expression and proteins were evaluated in the same cohort of animals that underwent behavioral testing at 3-, 6-, and 9-months of age, and the age at brain analysis was approximately 11-months of age. With consideration to the sexual dimorphism in prodromal AD and alcohol consumption in adolescence, rats of both sexes were utilized. Overall, the key findings are that the AD genotype accelerates AIE-dependent cognitive decline in female rats at 6 months of age, but male AD-rats are impaired on spatial navigation by 3-months with no additional deficits due to AIE exposure.

It was revealed that AD transgenes had unique effects on both neurotrophin-related and cell pathology marker proteins. Sex-specific effects were found as a function of AD transgenes: In males, AD transgenes raised pTau in the ventral hippocampus and decreased mNGF levels in the ventral hippocampus. Whereas in females, AD transgenes reduced vAChT levels and decreased proBDNF levels in the dorsal hippocampus. Additionally, while AD transgenes increased p75NTR levels, the impact varied by sex and brain region—female TgF344-AD rats exhibited increased p75NTR in the dorsal hippocampus, whereas male TgF344-AD rats displayed increased p75NTR levels in the ventral hippocampus.

Adolescent ethanol exposure had distinct effects on various factors. Both male and female rats exposed to AIE exhibited reduced TrkA levels in the dorsal hippocampus and selectively in the ventral hippocampus for males. In male rats, AIE also influenced pro-NGF levels in the dorsal hippocampus (decreasing) and ventral hippocampus (increasing), as well as increased pTau in the dorsal hippocampus.

Sex-specific interactions were observed between AD transgenes and AIE in the ventral hippocampus, with female rats showing decreased mNGF levels in the ventral hippocampus, while male rats exhibited increased proNGF, but reduced pro-BDNF in the ventral hippocampus due to AIE. Furthermore, male rats with AD transgenes had an increase in p75NTR. These sex-specific alterations do alter the ratio of immature to mature neurotrophins, leaning towards an increase in proneurotrophins over mature neurotrophins.

### 4.1. Sex influences the behavioral consequences of developmental ethanol exposure and AD transgenes

Female rats exposed to adolescent ethanol had an accelerated onset of spatial navigation impairments associated with AD transgenes: AIE-induced deficits on spontaneous alternation arise at 9-months in the WT female rats and at 6-months in the AD female rats. Male rats with AD transgenes had an earlier onset of impaired spatial navigation, at 3-months old, with no additional deficit due to AIE. At 6-months of age, both AD and AIE-exposed male rats were performing worse than WT-CON males at 3-months of age, regardless of Genotype or Exposure. Most male rats (56%) at 9 months of age did not meet the arm entry criteria on the plus maze.

This is in line with previous studies reporting reduced locomotor activity in F344 male rats as they age (Ramos et al., 1997; Rex et al., 1999). Thus, full conclusions about the spatial navigation decline across adulthood in the male TgF344-AD rats was not determined. Previous studies have suggested an earlier age-related decline in spatial memory in wildtype male rats, when compared to female rats, in both F344 (Febo et al., 2020) and Sprague Dawley strains (Reitz et al., 2021). The AD genotype may be accelerating the natural age-related cognitive decline seen in males to as early as 3-months old. Short-term spatial memory performance on the DNMTP task was somewhat inconsistent across training days and delay intervals. However, there were clear sex-effects: In female rats, both AD transgenes and adolescent ethanol exposure impaired short-term memory performance. In contrast, the opposite effect was found in male rats: AIE and AD transgenes improved short-term memory performance. Another study found that TgF344-AD rats develop strong preference for one of the two levers, that prevented memory assessment with DMTP in 50% of rats (Sagalajev et al., 2023). In AD-rats that did not develop lever-preferences, and no spatial short-term memory impairments were observed on either DMTP (Sagalajev et al., 2023) or DNMTP (Muñoz-Moreno et al., 2018). Those studies did not include sex as a factor, which we show can influence this behavioral assay.

Social recognition memory is thought to largely be dependent on the ventral hippocampus (Bannerman et al., 2004; Okuyama et al., 2016), an area that is often targeted by amyloid plaque accumulation and AIE-induced brain pathology (Galaj et al., 2019; Russo et al., 2021; Tsui et al., 2022a). Animals were initially tested on social recognition at 3-months of age, only one month after the completion of AIE. Research suggests that rodents that experience chronic diminished conditions tend to be more sensitive to the initial stress and anxiety caused by behavioral testing (Foster, 2023). There is concern that there could be carryover effects from previous behavioral testing due to the longitudinal design with multiple behavioral tasks (Cnops et al, 2022). This concern needs to be balanced with the benefits of longitudinal aging studies that can better identify the onset of cognitive impairment and control for cohort effects.

Given concerns that locomotor activity declines with age in rodents (Altun et al., 2007; Casadesus et al., 2001), and the decrease in arm entries we saw during spontaneous alternation, all rats were tested for general activity at 11-months of age. Although we did not statistically compare across sexes, male rats appear to move less than female rats in the activity chambers. When separated by sex, AD female rats moved less than wild-type female rats, while male rats did not differ by Exposure or Genotype. Exposure to AIE did not influence movement in the chamber, suggesting that the acceleration of memory impairments in the AD rats exposed to AIE is not due to a lack of activity during the memory tasks. However, female rats with AD transgenes have been shown to have heightened anxiety scores (Saré et al., 2020; Srivastava et al., 2023), which is suggested by the decreased time spent in the center of the activity chambers in AD female rats; although, female rats with AD transgenes exposed to adolescent ethanol did not display this suppression of activity in the center. Thus, anxiety-like behavior likely did not contribute to the deficits in spatial navigation observed in these rats.

### 4.2. Sex-specific changes in AD-pathological markers as a function of developmental ethanol exposure

Although neurotrophin levels have not been published in the Tg344-AD model, other AD rodent models (mouse 5xFAD, McGill-R-Thy1-APP rat) exhibit dysregulation of pro- and mature forms of NGF and BDNF. Protein levels of proNGF are increased in the cortex of McGill-R-Thy1-APP rat, compared to wild-type rats, were as mRNA for NGF is unchanged. In contrast, there was no change in protein levels of proBDNF, but mRNA for mBDNF decreased in the McGill-R-Thy1-APP model (Iulita et al., 2017). In the mouse 5xFAD model, BDNF is also reduced in the dorsal hippocampus, but NGF and the corresponding pro-neurotrophins (proNGF, proBDNF) were not measured (de Pins et al., 2019). However, these previous studies did not include sex as a variable.

Protein levels of pathological markers and neurotrophin-related factors and were assessed across the two main hippocampal sectors to reveal the neurobiological underpinnings of the cognitive impairments associated with AD transgenes and adolescent ethanol exposure as a function of sex. The key findings are the pathological markers of pTau-181 and VAChT, which modulates ACh release, are modulated in a sex-specific manner. Male rats, but not female rats, exposed to AIE had an increase in pTau-181 in the dorsal hippocampus, and AD transgenes selectively in males led to increased pTau-181 in the ventral hippocampus. In female rats, AD transgenes suppressed VAChT in the dorsal hippocampus, with male rats having no changes in VAChT. Markers of both pTau (Mendes et al, 2024) and PET imaging of VAChT (Aghourian et al., 2017) are used as markers to quantify AD severity, and our results demonstrate that sex appears to influence which is an effective marker for AD-related pathology.

We selected several neurotropic factors to assess, as our original hypothesis was that changes the pro- to mature neurotrophin balance, as well as the Trk (tropomyosin-receptor-kinase)/p75 NTR balance, could lead to sex-specific vulnerabilities following adolescent intermittent ethanol exposure in the AD-predisposed brain. Reductions in TrkA, or an increase in p75NTR, promotes proNGF to activate p75NTR-dependent apoptotic pathways (Fahnestock & Shekari, 2019). Studies suggest that alterations in TrkA signaling may contribute to the neurodegenerative processes observed in AD. TrkA is increasingly lost from mild cognitive impairment (MCI) to AD (Fahnestock & Shekari, 2019; Mufson et al., 2019). Dysregulation of p75NTR signaling is linked to neurodegeneration and may contribute to the progression of AD (Boissière et al., 1996). Investigating the intricate balance between neurotrophins TrkA and p75NTR will provide valuable insights into the molecular mechanisms underlying the susceptibility and progression AD, offering potential avenues for therapeutic interventions.

The present study found that AD transgenes and AIE exposure altered levels of pro- and mNGF and their respective receptors. Female rats with AD transgenes that have been exposed to AIE had a decrease in mNGF in the vHPC (not measured in dHPC). Whereas in male rats, AD transgenes resulted in decreased mNGF and increased proNGF levels in the vHPC, but not dHPC. Furthermore, AD transgenes increased p75NTR levels, but the impact varied by sex and brain region: Female AD rats exhibited increased p75NTR levels in the dHPC, whereas male AD rats displayed increased p75NTR levels in the vHPC. Additionally, AIE decreased TrkA in the dHPC of both sexes, and also in the vHPC of male rats. Thus, there are several changes in pro- and mNGF levels, as well as TrkA and p75NTRs following AIE and aging with AD transgenes. Such changes shift the balance from pro-survival to apoptosis and neurite retraction (Fahnestock & Shekari, 2019). Thus, AIE-induced suppression of mature NTs and altered TrkA/p75NTR ratio creates a brain that is vulnerable to age-related and AD-related pathologies, which leads to an acceleration of cognitive decline, particularly in female rats.

The role of NGF/TrkA and proNGF/p75NTR in maintaining basal forebrain cholinergic neurons is well established (Cuello et al., 2010; Iulita et al., 2017). It has been demonstrated that Aβ_1−42_binding to p75^NTR^ leads to an increase in the extracellular levels of Aβ_1−42_, which then causes neurotoxicity and apoptosis (Yaar et al., 1997) and further meditates Aβ-induced tau hyperphosphorylation (Shen et al., 2019). pTau protein expression was increased in AIE-exposed male rats, which suggests that pTau may have an earlier onset in males with AD transgenes if they were exposed to heavy ethanol exposure during adolescence. This is pertinent as neurofibrillary tangles in basal forebrain cholinergic neurons have been found in patients with MCI, and neurofibrillary tangles burden in the basal forebrain correlates with memory decline (Mesulam et al., 2004). Tau phosphorylation can cause dysregulation of axonal transport, including the retrograde transport of NGF from the hippocampus to the MS/DB through cholinergic neurons (Butler et al., 2019; Combs et al., 2019). Miscommunication between the cholinergic signaling within the septohippocampal pathway may cause an array of downstream effects including interferences with neurogenesis, heightened inflammation, and cognitive decline (Anacker & Hen, 2017; Clelland et al., 2009; Macht et al., 2020; Macht et al., 2022).

While NGF is critical to the survival and maintenance of cholinergic neurons in the basal forebrain, BDNF is critical for the maintenance of entorhinal cortical projection neurons to the hippocampus (Falkenberg et al., 1993; Roysommuti & Wyss, 2022). BDNF is anterogradely transported from production sites in the ERC to the hippocampus, and has been established as a modulator of synaptic plasticity in the hippocampus (Gooney & Lynch, 2001; Kang & Schuman, 1995; Lu, 2003). In patients with AD, as well as in rodent models of AD, BDNF and its neurotrophic receptor TrkB are downregulated (Ginsberg et al., 2019; Hock et al., 2000; Tanila, 2017). Projection neurons from the ERC, through BDNF, heavily contribute to the initial memory impairments seen in early-stage AD which makes them an especially timely target to early intervention for AD (Erickson et al., 2010; Nagahara et al., 2009; Roysommuti & Wyss, 2022; Zhang et al., 2022). Interestingly, levels of mBDNF remained unchanged by AD transgenes or adolescent ethanol exposure across the hippocampus of both male and female rats. In contrast, levels of proBDNF protein expression in the dorsal hippocampus were decreased in AD female rats, but were slightly increased in female rats exposed to AIE. In male rats with AD transgenes, adolescent ethanol exposure decreased proBDNF protein expression in the ventral hippocampus. There is also a reduction in proBDNF in human patients with MCI (Mufson et al., 2007), which get larger as patients transition into AD (Peng et al., 2005). Such changes may be compensatory to the early and middle stages of neurodegeneration, but fail as the disease progresses.

### 4.3 Conclusions

There were unique sex-dependent effects of AD transgenes, adolescent ethanol exposure, and the interaction of those variables on behavior and pathological and neurotrophin markers.

Male rats with AD transgenes display spatial location impairment at 3 months of age, and this was not amplified by adolescent ethanol exposure. Pathological changes were observed in male rats as a function of AD transgenes and AIE: There was an increase in pTau and there was a shift in the balance of proNGF/NGF and its receptors toward cell death mechanisms.

There was an acceleration of AD-induced cognitive decline in TgF344-AD rats due to exposure to AIE in female rats. A reduction in TrkA receptors due to AIE, and a decrease in VAChT due to AD transgenes, were found within the dorsal hippocampus of female rats. A dysfunctional cholinergic system may be a significant factor in the accelerated spatial navigation impairments in females with AD transgenes exposed to adolescent ethanol.

This study is one of the few that examined whether sex differentially contributed to the progression of behavioral impairments associated with AD transgenes with and without developmental alcohol exposure. Although the longitudinal design allowed for tracking of behavioral changes across time, it did not permit for examination of the brain at different time points. Thus, we cannot fully determine which changes in the brain at 11 months track with the onset of cognitive dysfunction. A cross sectional analysis of brain pathology is needed.

## Author contributions

**Nicole Reitz**: Conceptualization, Methodology, Investigation, Writing-Original draft preparation, Formal Analysis, Visualization; **Polliana Nunes**: Methodology, Investigation, Writing-Reviewing and Editing, Formal Analysis, Visualization; **Lisa Savage**: Conceptualization, Methodology, Writing-Original draft preparation; Supervision, Validation, Funding acquisition

## List of Abbreviations

Aβ: amyloid beta
AD: Alzheimer’s Disease
APP: amyloid precursor protein
BDNF: brain-derived neurotrophic factor
BEC: blood ethanol content
ChAT: choline acetyltransferase
CON: water control exposure
DNMTP: delayed nonmatching to position
PSEN: presenilin
AIE: adolescent intermittent ethanol exposure
MS/DB: medial septum/diagonal band
NOP: novel object location/place
NGF: nerve growth factor
pTau: phosphorylated tau
TrkA: tropomyosin receptor kinase A
TrKB: tropomyosin receptor kinase B
WT: Wildtype
VAChT: vesicular acetylcholine transporter

## References

1. Aghourian M, Legault-Denis C, Soucy JP, Rosa-Neto P, Gauthier S, Kostikov A, et al. Quantification of brain cholinergic denervation in Alzheimer’s disease using PET imaging with [18F]-FEOBV. Mol Psychiatry. 2017;22(11):1531–8. doi: 10.1038/mp.2017.183

2. Altun M, Bergman E, Edström E, Johnson H, Ulfhake B. Behavioral impairments of the aging rat. Physiol Behav. 2007;92(5):911–23. doi: 10.1016/j.physbeh.2007.06.017

3. Anacker C, Hen R. Adult hippocampal neurogenesis and cognitive flexibility - linking memory and mood. Nat Rev Neurosci. 2017;18(6):335–46. doi: 10.1038/nrn.2017.45

4. Bannerman DM, Rawlins JNP, McHugh SB, Deacon RMJ, Yee BK, Bast T, et al. Regional dissociations within the hippocampus--memory and anxiety. Neurosci Biobehav Rev. 2004;28(3):273–83. doi: 10.1016/j.neubiorev.2004.03.004

5. Barnett A, David E, Rohlman A, Nikolova VD, Moy SS, Vetreno RP, et al. Adolescent Binge Alcohol Enhances Early Alzheimer’s Disease Pathology in Adulthood Through Proinflammatory Neuroimmune Activation. Front Pharmacol. 2022;13:884170. Doi: 10.3389/fphar.2022.884170

6. Berkowitz LE, Harvey RE, Drake E, Thompson SM, Clark BJ. Progressive impairment of directional and spatially precise trajectories by TgF344-Alzheimer’s disease rats in the Morris Water Task. Sci Rep. 2018;8:16153. Doi: 10.1038/s41598-018-34368-w

7. Bert B, Fink H, Huston JP, Voits M. Fischer 344 and wistar rats differ in anxiety and habituation but not in water maze performance. Neurobiol Learn Mem. 2002;78(1):11–22. Doi: 10.1006/nlme.2001.4040

8. Boissière F, Lehéricy S, Strada O, Agid Y, Hirsch EC. Neurotrophin receptors and selective loss of cholinergic neurons in Alzheimer disease. Mol Chem Neuropathol. 1996;28(1–3):219–23. Doi: 10.1007/BF02815225

9. Butler VJ, Salazar DA, Soriano-Castell D, Alves-Ferreira M, Dennissen FJA, Vohra M, et al. Tau/MAPT disease-associated variant A152T alters tau function and toxicity via impaired retrograde axonal transport. Hum Mol Genet. 2019;28(9):1498–514. Doi: 10.1093/hmg/ddy442

10. Casadesus G, Shukitt-Hale B, Joseph JA. Automated measurement of age-related changes in the locomotor response to environmental novelty and home-cage activity. Mech Ageing Dev. 2001;122(15):1887–97. Doi: 10.1016/s0047-6374(01)00324-4

11. Chaney AM, Lopez-Picon FR, Serrière S, Wang R, Bochicchio D, Webb SD, et al. Prodromal neuroinflammatory, cholinergic and metabolite dysfunction detected by PET and MRS in the TgF344-AD transgenic rat model of AD: a collaborative multi-modal study. Theranostics. 2021;11(14):6644–67. Doi: 10.7150/thno.56059

12. Clelland CD, Choi M, Romberg C, Clemenson GD, Fragniere A, Tyers P, et al. A functional role for adult hippocampal neurogenesis in spatial pattern separation. Science. 2009;325(5937):210–3. Doi: 10.1126/science.1173215

13. Cohen RM, Rezai-Zadeh K, Weitz TM, Rentsendorj A, Gate D, Spivak I, et al. A transgenic Alzheimer rat with plaques, tau pathology, behavioral impairment, oligomeric aβ, and frank neuronal loss. J Neurosci Off J Soc Neurosci. 2013;33(15):6245–56. Doi: 10.1523/JNEUROSCI.3672-12.2013

14. Coleman LG, Crews FT, Vetreno RP. The persistent impact of adolescent binge alcohol on adult brain structural, cellular, and behavioral pathology: A role for the neuroimmune system and epigenetics. Int Rev Neurobiol. 2021;160:1–44. Doi: 10.1016/bs.irn.2021.08.001

15. Coleman LG, He J, Lee J, Styner M, Crews FT. Adolescent binge drinking alters adult brain neurotransmitter gene expression, behavior, brain regional volumes, and neurochemistry in mice. Alcohol Clin Exp Res. 2011;35(4):671–88. Doi: 10.1111/j.1530-0277.2010.01385.x

16. Combs B, Mueller RL, Morfini G, Brady ST, Kanaan NM. Tau and Axonal Transport Misregulation in Tauopathies. Adv Exp Med Biol. 2019;1184:81–95. Doi: 10.1007/978-981-32-9358-8_7

17. Cnops V, Iyer VR, Parathy N, Wong P, Dawe GS. Test, rinse repeat: A review of carryover effects in rodent behavioral assays. Neurosci Biobehav Rev. 2022;135:104560. Doi: 10.1016

18. Cuello AC, Bruno MA, Allard S, Leon W, Iulita MF. Cholinergic involvement in Alzheimer’s disease. A link with NGF maturation and degradation. J Mol Neurosci MN. 2010;40(1– 2):230–5. Doi: 10.1007/s12031-009-9238-z

19. Cuello AC, Pentz R, Hall H. The Brain NGF Metabolic Pathway in Health and in Alzheimer’s Pathology. Front Neurosci. 2019;13:62. Doi: 10.3389/fnins.2019.00062

20. Cuello AC, Bruno MA, Bell KFS. NGF-cholinergic dependency in brain aging, MCI and Alzheimer’s disease. Curr Alzheimer Res. 2007;4(4):351–8. Doi: 10.2174/156720507781788774

21. de Pins B, Cifuentes-Díaz C, Farah AT, López-Molina L, Montalban E, Sancho-Balsells A, et al. Conditional BDNF Delivery from Astrocytes Rescues Memory Deficits, Spine Density, and Synaptic Properties in the 5xFAD Mouse Model of Alzheimer Disease. J Neurosci Off J Soc Neurosci. 2019;39(13):2441–58. Doi: 10.1523/JNEUROSCI.2121-18.2019

22. Drummond E, Wisniewski T. Alzheimer’s disease: experimental models and reality. Acta Neuropathol (Berl). 2017 Feb;133(2):155–75. Doi: 10.1007/s00401-016-1662-x

23. Erickson KI, Prakash RS, Voss MW, Chaddock L, Heo S, McLaren M, et al. Brain-Derived Neurotrophic Factor Is Associated with Age-Related Decline in Hippocampal Volume. J Neurosci. 2010;30(15):5368–75. Doi: 10.1523/JNEUROSCI.6251-09.2010

24. Fahnestock M, Shekari A. ProNGF and Neurodegeneration in Alzheimer’s Disease. Front Neurosci. 2019;13:129. Doi: 10.3389/fnins.2019.00129

25. Falkenberg T, Metsis M, Timmusk T, Lindefors N. Entorhinal cortex regulation of multiple brain-derived neurotrophic factor promoters in the rat hippocampus. Neuroscience. 1993 Dec;57(4):891–6. Doi: 10.1016/0306-4522(93)90034-d

26. Febo M, Rani A, Yegla B, Barter J, Kumar A, Wolff CA, et al. Longitudinal Characterization and Biomarkers of Age and Sex Differences in the Decline of Spatial Memory. Front Aging Neurosci. 2020;12:34. Doi: 10.3389/fnagi.2020.00034

27. Fernandez GM, Savage LM. Adolescent binge ethanol exposure alters specific forebrain cholinergic cell populations and leads to selective functional deficits in the prefrontal cortex. Neuroscience. 2017;361:129–43. Doi: 10.1016/j.neuroscience.2017.08.013

28. Foster TC. Animal models for studies of alcohol effects on the trajectory of age-related cognitive decline. Alcohol. 2023;107:4–11. Doi: 10.1016/j.alcohol.2022.04.005

29. Galaj E, Kipp BT, Floresco SB, Savage LM. Persistent Alterations of Accumbal Cholinergic Interneurons and Cognitive Dysfunction after Adolescent Intermittent Ethanol Exposure. Neuroscience. 2019;404:153–64. doi: 10.1016/j.neuroscience.2019.01.062

30. Galloway CR, Ravipati K, Singh S, Lebois EP, Cohen RM, Levey AI, et al. Hippocampal place cell dysfunction and the effects of muscarinic M1 receptor agonism in a rat model of Alzheimer’s disease. Hippocampus. 2018;28(8):568–85. doi: 10.1002/hipo.22961

31. Gilmor ML, Erickson JD, Varoqui H, Hersh LB, Bennett DA, Cochran EJ, et al. Preservation of nucleus basalis neurons containing choline acetyltransferase and the vesicular acetylcholine transporter in the elderly with mild cognitive impairment and early Alzheimer’s disease. J Comp Neurol. 1999;411(4):693–704.

32. Ginsberg SD, Malek-Ahmadi MH, Alldred MJ, Che S, Elarova I, Chen Y, et al. Selective decline of neurotrophin and neurotrophin receptor genes within CA1 pyramidal neurons and hippocampus proper: correlation with cognitive performance and neuropathology in mild cognitive impairment and Alzheimer’s disease. Hippocampus. 2019;29(5):422–39. doi: 10.1002/hipo.22802

33. Ginsberg SD, Malek-Ahmadi MH, Alldred MJ, Chen Y, Chen K, Chao MV, et al. Brain-derived neurotrophic factor (BDNF) and TrkB hippocampal gene expression are putative predictors of neuritic plaque and neurofibrillary tangle pathology. Neurobiol Dis. 2019;132:104540. doi: 10.1016/j.nbd.2019.104540

34. Gooney M, Lynch MA. Long-term potentiation in the dentate gyrus of the rat hippocampus is accompanied by brain-derived neurotrophic factor-induced activation of TrkB. J Neurochem. 2001;77(5):1198–207. doi: 10.1046/j.1471-4159.2001.00334.x

35. Hagg T, Fass-Holmes B, Vahlsing HL, Manthorpe M, Conner JM, Varon S. Nerve growth factor (NGF) reverses axotomy-induced decreases in choline acetyltransferase, NGF receptor and size of medial septum cholinergic neurons. Brain Res. 1989;505(1):29–38. doi: 10.1016/0006-8993(89)90112-1

36. Hagg T, Manthorpe M, Vahlsing HL, Varon S. Delayed treatment with nerve growth factor reverses the apparent loss of cholinergic neurons after acute brain damage. Exp Neurol. 1988;101(2):303–12. doi: 10.1016/0014-4886(88)90013-1

37. Hock C, Heese K, Hulette C, Rosenberg C, Otten U. Region-specific neurotrophin imbalances in Alzheimer disease: decreased levels of brain-derived neurotrophic factor and increased levels of nerve growth factor in hippocampus and cortical areas. Arch Neurol. 2000;5(6):846–51. doi: 10.1001/archneur.57.6.846

38. Holland D, Desikan RS, Dale AM, McEvoy LK. Higher Rates of Decline for Women and Apolipoprotein E ε4 Carriers. AJNR Am J Neuroradiol. 2013;34(12):2287–93. doi: 10.3174/ajnr.A3601

39. Iulita MF, Bistué Millón MB, Pentz R, Aguilar LF, Do Carmo S, Allard S, et al. Differential deregulation of NGF and BDNF neurotrophins in a transgenic rat model of Alzheimer’s disease. Neurobiol Dis. 2017;108:307–23. doi: 10.1016/j.nbd.2017.08.019

40. Kang H, Schuman EM. Long-lasting neurotrophin-induced enhancement of synaptic transmission in the adult hippocampus. Science. 1995;267(5204):1658–62. doi: 10.1126/science.7886457

41. Kipp BT, Nunes PT, Galaj E, Hitchcock B, Nasra T, Poynor KR, et al. Adolescent Ethanol Exposure Alters Cholinergic Function and Apical Dendritic Branching Within the Orbital Frontal Cortex. Neuroscience. 2021;473:52–65. doi: 10.1016/j.neuroscience.2021.08.014

42. Ledesma JC, Rodríguez-Arias M, Gavito AL, Sánchez-Pérez AM, Viña J, Medina Vera D, et al. Adolescent binge-ethanol accelerates cognitive impairment and β-amyloid production and dysregulates endocannabinoid signaling in the hippocampus of APP/PSE mice. Addict Biol. 2021;26(1):e12883. doi: 10.1111/adb.12883

43. León BE, Kang S, Franca-Solomon G, Shang P, Choi DS. Alcohol-Induced Neuroinflammatory Response and Mitochondrial Dysfunction on Aging and Alzheimer’s Disease. Front Behav Neurosci. 2021;15:778456. doi: 10.3389/fnbeh.2021.778456

44. Lin KA, Choudhury KR, Rathakrishnan BG, Marks DM, Petrella JR, Doraiswamy PM, et al. Marked gender differences in progression of mild cognitive impairment over 8 years. Alzheimers Dement N Y N. 2015;1(2):103–10. doi: 10.1016/j.trci.2015.07.001

45. Lu B. BDNF and activity-dependent synaptic modulation. Learn Mem Cold Spring Harb N. 2003;10(2):86–98. doi: 10.1101/lm.54603

46. Macht VA, Crews FT, Vetreno RP. Neuroimmune and epigenetic mechanisms underlying persistent loss of hippocampal neurogenesis following adolescent intermittent ethanol exposure. Curr Opin Pharmacol. 2020;50:9–16. doi: 10.1016/j.coph.2019.10.007

47. Macht VA, Vetreno RP, Crews FT. Cholinergic and Neuroimmune Signaling Interact to Impact Adult Hippocampal Neurogenesis and Alcohol Pathology Across Development. Front Pharmacol. 2022;13:849997. doi: 10.3389/fphar.2022.849997

48. Mathiasen JR, DiCamillo A. Social recognition assay in the rat. Curr Protoc Neurosci. 2010;Chapter 8:Unit 8.5I. doi: doi.org/10.1002/0471142301.ns0805is53

49. Maynard ME, Barton EA, Robinson CR, Wooden JI, Leasure JL. Sex differences in hippocampal damage, cognitive impairment, and trophic factor expression in an animal model of an alcohol use disorder. Brain Struct Funct. 2018;223(1):195–210. doi: 10.1007/s00429-017-1482-3

50. Mesulam M, Shaw P, Mash D, Weintraub S. Cholinergic nucleus basalis tauopathy emerges early in the aging-MCI-AD continuum. Ann Neurol. 2004;55(6):815–28. doi: 10.1002/ana.20100

51. Mielke MM. Sex and Gender Differences in Alzheimer’s Disease Dementia. Psychiatr Times. 2018;35(11):14–7. PMC6390276

52. Mufson EJ, Counts SE, Fahnestock M, Ginsberg SD. Cholinotrophic molecular substrates of mild cognitive impairment in the elderly. Curr Alzheimer Res. 2007;4(4):340–50. doi: 10.2174/156720507781788855

53. Mufson EJ, Counts SE, Ginsberg SD, Mahady L, Perez SE, Massa SM, et al. Nerve Growth Factor Pathobiology During the Progression of Alzheimer’s Disease. Front Neurosci. 2019;13:533. doi: 10.3389/fnins.2019.00533

54. Muñoz-Moreno E, Tudela R, López-Gil X, Soria G. Early brain connectivity alterations and cognitive impairment in a rat model of Alzheimer’s disease. Alzheimers Res Ther. 2018;10(1):16. doi: 10.1186/s13195-018-0346-2

55. Nagahara AH, Merrill DA, Coppola G, Tsukada S, Schroeder BE, Shaked GM, et al. Neuroprotective effects of brain-derived neurotrophic factor in rodent and primate models of Alzheimer’s disease. Nat Med. 2009;15(3):331–7. doi: doi.org/10.1038/nm.1912

56. Okuyama T, Kitamura T, Roy DS, Itohara S, Tonegawa S. Ventral CA1 neurons store social memory. Science. 2016;353(6307):1536–41. doi: 10.1126/science.aaf7003

57. Parent MJ, Bedard MA, Aliaga A, Minuzzi L, Mechawar N, Soucy JP, et al. Cholinergic Depletion in Alzheimer’s Disease Shown by [(18) F]FEOBV Autoradiography. Int J Mol Imaging. 2013;2013:205045. doi: 10.1155/2013/205045

58. Peng S, Wuu J, Mufson EJ, Fahnestock M. Precursor form of brain-derived neurotrophic factor and mature brain-derived neurotrophic factor are decreased in the pre-clinical stages of Alzheimer’s disease. J Neurochem. 2005;93(6):1412–21. doi: 10.1111/j.1471-4159.2005.03135.x

59. Ramos A, Berton O, Mormède P, Chaouloff F. A multiple-test study of anxiety-related behaviours in six inbred rat strains. Behav Brain Res. 1997 Apr;85(1):57–69. doi: 10.1016/s0166-4328(96)00164-7

60. Reitz NL, Nunes PT, Savage LM. Adolescent Binge-Type Ethanol Exposure in Rats Mirrors Age-Related Cognitive Decline by Suppressing Cholinergic Tone and Hippocampal Neurogenesis. Front Behav Neurosci. 2021;15:772857. doi: 10.3389/fnbeh.2021.772857

61. Rex A, Sondern U, Voigt JP, Franck S, Fink H. Strain differences in fear-motivated behavior of rats. Pharmacol Biochem Behav. 1996;54(1):107–11. doi: 10.1016/0091-3057(95)02128-0

62. Rex A, Voigt JP, Fink H. Behavioral and neurochemical differences between Fischer 344 and Harlan-Wistar rats raised identically. Behav Genet. 1999;29(3):187–92. doi: 10.1023/a:1021644002588

63. Rorabaugh JM, Chalermpalanupap T, Botz-Zapp CA, Fu VM, Lembeck NA, Cohen RM, et al. Chemogenetic locus coeruleus activation restores reversal learning in a rat model of Alzheimer’s disease. Brain J Neurol. 2017;140(11):3023–38. doi: 10.1093/brain/awx232

64. Roysommuti S, Wyss JM. Brain-Derived Neurotrophic Factor Potentiates Entorhinal-Dentate but not Hippocampus CA1 Pathway in Adult Male Rats: A Mechanism of Taurine-Modulated BDNF on Learning and Memory. Adv Exp Med Biol. 2022;1370:369–79. doi: 10.1007/978-3-030-93337-1_35

65. Russo ML, Molina-Campos E, Ybarra N, Rogalsky AE, Musial TF, Jimenez V, et al. Variability in sub-threshold signaling linked to Alzheimer’s disease emerges with age and amyloid plaque deposition in mouse ventral CA1 pyramidal neurons. Neurobiol Aging. 2021;106:207–22. doi: 10.1016/j.neurobiolaging.2021.06.018

66. Sagalajev B, Lennartz L, Vieth L, Gunawan CT, Neumaier B, Drzezga A, et al. TgF344-AD Rat Model of Alzheimer’s Disease: Spatial Disorientation and Asymmetry in Hemispheric Neurodegeneration. J Alzheimers Dis Rep. 2023;7(1):1085–94. doi: 10.3233/ADR-230038

67. Sanna PP, Cabrelle C, Kawamura T, Mercatelli D, O’Connor N, Roberts AJ, et al. A History of Repeated Alcohol Intoxication Promotes Cognitive Impairment and Gene Expression Signatures of Disease Progression in the 3xTg Mouse Model of Alzheimer’s Disease. eNeuro. 2023;10(7):ENEURO.0456-22.2023. doi: 10.1523/ENEURO.0456-22.2023

68. Saré RM, Cooke SK, Krych L, Zerfas PM, Cohen RM, Smith CB. Behavioral Phenotype in the TgF344-AD Rat Model of Alzheimer’s Disease. Front Neurosci. 2020;14:601. doi: 10.3389/fnins.2020.00601

69. Scott SA, Crutcher KA. Nerve growth factor and Alzheimer’s disease. Rev Neurosci. 1994;5(3):179–211. doi: 10.1515/revneuro.1994.5.3.179

70. Shen LL, Li WW, Xu YL, Gao SH, Xu MY, Bu XL, et al. Neurotrophin receptor p75 mediates amyloid β-induced tau pathology. Neurobiol Dis. 2019;132:104567. doi: 10.1016/j.nbd.2019.104567

71. Spear LP. Effects of adolescent alcohol consumption on the brain and behaviour. Nat Rev Neurosci. 2018;19(4):197–214. doi: 10.1038/nrn.2018.10

72. Srivastava H, Lasher AT, Nagarajan A, Sun LY. Sexual dimorphism in the peripheral metabolic homeostasis and behavior in the TgF344-AD rat model of Alzheimer’s disease. Aging Cell. 2023;e13854. doi: 10.1111/acel.13854

73. Tanila H. The role of BDNF in Alzheimer’s disease. Neurobiol Dis. 2017;97(Pt B):114–8. doi: 10.1016/j.nbd.2016.05.008

74. Tsui KC, Roy J, Chau SC, Wong KH, Shi L, Poon CH, et al. Distribution and inter-regional relationship of amyloid-beta plaque deposition in a 5xFAD mouse model of Alzheimer’s disease. Front Aging Neurosci. 2022;14:964336. doi: 10.3389/fnagi.2022.964336

75. Tucker AE, Alicea Pauneto CDM, Barnett AM, Coleman LG. Chronic Ethanol Causes Persistent Increases in Alzheimer’s Tau Pathology in Female 3xTg-AD Mice: A Potential Role for Lysosomal Impairment. Front Behav Neurosci. 2022;16:886634. doi: 10.3389/fnbeh.2022.886634

76. Vetreno RP, Bohnsack JP, Kusumo H, Liu W, Pandey SC, Crews FT. Neuroimmune and epigenetic involvement in adolescent binge ethanol-induced loss of basal forebrain cholinergic neurons: Restoration with voluntary exercise. Addict Biol. 2020;25(2):e12731. doi: 10.1111/adb.12731

77. Vetreno RP, Broadwater M, Liu W, Spear LP, Crews FT. Adolescent, but Not Adult, Binge Ethanol Exposure Leads to Persistent Global Reductions of Choline Acetyltransferase Expressing Neurons in Brain. PLoS ONE. 2014;9(11):e113421. doi: 10.1371/journal.pone.0113421

78. Vetreno RP, Lawrimore CJ, Rowsey PJ, Crews FT. Persistent Adult Neuroimmune Activation and Loss of Hippocampal Neurogenesis Following Adolescent Ethanol Exposure: Blockade by Exercise and the Anti-inflammatory Drug Indomethacin. Front Neurosci. 2018;12:200. doi: 10.3389/fnins.2018.00200

79. Voorhees JR, Remy MT, Erickson CM, Dutca LM, Brat DJ, Pieper AA. Occupational-like organophosphate exposure disrupts microglia and accelerates deficits in a rat model of Alzheimer’s disease. NPJ Aging Mech Dis. 2019;5:3. doi: 10.1038/s41514-018-0033-3

80. Wu C, Yang L, Li Y, Dong Y, Yang B, Tucker LD, et al. Effects of Exercise Training on Anxious-Depressive-like Behavior in Alzheimer Rat. Med Sci Sports Exerc. 2020;52(7):1456–69. doi: 10.1249/MSS.0000000000002294

81. Yaar M, Zhai S, Pilch PF, Doyle SM, Eisenhauer PB, Fine RE, et al. Binding of beta-amyloid to the p75 neurotrophin receptor induces apoptosis. A possible mechanism for Alzheimer’s disease. J Clin Invest. 1997;100(9):2333–40. doi: 10.1172/JCI119772

82. Zhang W, Ge MM, Zhang LQ, Yuan XM, Han SY, Manyande A, et al. Dysfunction of the Brain-derived Neurotrophic Factor-Tyrosine Kinase B Signaling Pathway Contributes to Learning and Memory Impairments Induced by Neuroinflammation in Mice. Neuroscience. 2022;505:21–33. doi: 10.1016/j.neuroscience.2022.10.003

